# Programmable protein degraders enable selective knockdown of pathogenic β-catenin subpopulations *in vitro* and *in vivo*

**DOI:** 10.1101/2024.11.10.622803

**Authors:** Tianzheng Ye, Azmain Alamgir, Cara M. Robertus, Darianna Colina, Connor Monticello, Thomas Connor Donahue, Lauren Hong, Sophia Vincoff, Shrey Goel, Peter Fekkes, Luis Miguel Camargo, Kieu Lam, James Heyes, David Putnam, Christopher A. Alabi, Pranam Chatterjee, Matthew P. DeLisa

**Author notes:** To whom correspondence should be addressed: Pranam Chatterjee, Department of Biomedical Engineering, Duke University, Durham, NC 27708 USA; Matthew P. DeLisa, Robert F. Smith School of Chemical and Biomolecular Engineering, Cornell University, Ithaca, NY 14853 USA.

## Abstract

Aberrant activation of Wnt signaling results in unregulated accumulation of cytosolic β-catenin, which subsequently enters the nucleus and promotes transcription of genes that contribute to cellular proliferation and malignancy. Here, we sought to eliminate pathogenic β-catenin from the cytosol using designer ubiquibodies (uAbs), chimeric proteins composed of an E3 ubiquitin ligase and a target-binding domain that redirect intracellular proteins to the proteasome for degradation. To accelerate uAb development, we leveraged a protein language model (pLM)-driven algorithm called SaLT&PepPr to computationally design “guide” peptides with affinity for β-catenin, which were subsequently fused to the catalytic domain of a human E3 called C-terminus of Hsp70-interacting protein (CHIP). Expression of the resulting peptide-guided uAbs in colorectal cancer cells led to the identification of several designs that significantly reduced the abnormally stable pool of free β-catenin in the cytosol and nucleus while preserving the normal membrane-associated subpopulation. This selective knockdown of pathogenic β-catenin suppressed Wnt/β-catenin signaling and impaired tumor cell survival and proliferation. Furthermore, one of the best degraders selectively decreased cytosolic but not membrane-associated β-catenin levels in livers of BALB/c mice following delivery as a lipid nanoparticle (LNP)-encapsulated mRNA. Collectively, these findings reveal the unique ability of uAbs to selectively eradicate abnormal proteins *in vitro* and *in vivo* and open the door to peptide-programmable biologic modulators of other disease-causing proteins.

## INTRODUCTION

The Wnt signaling pathway is a crucial regulator of various cellular processes, including cell proliferation, differentiation, and migration, and determination of cell fate during embryonic development and tissue homeostasis ^1-3^. A key downstream component of this pathway is β-catenin, a dual-function protein that plays an important role in cell-cell adhesion through interaction with E-cadherin and transcriptional regulation of Wnt-responsive genes through interaction with TCF/LEF transcription factors ^4,5^. Under normal conditions, β-catenin is kept in check by a destruction complex composed of several proteins including adenomatous polyposis coli (APC) and glycogen synthase kinase 3 (GSK3), which targets it for proteasomal degradation (**Fig. 1**). However, when Wnt signaling is aberrantly activated, such as from loss-of-function mutations in the *APC* gene or gain-of-function point mutations or deletions in *CTNNB1*, the gene encoding β-catenin, this degradation is blocked. Abnormally stabilized β-catenin accumulates in the cytoplasm and then translocates to the nucleus where it interacts with TCF/LEF to drive the expression of oncogenes such as *c-Myc* and *Cyclin D1* ^6-8^. This unchecked β-catenin activity promotes uncontrolled cell proliferation and survival, contributing to the development of various malignancies including colorectal cancer (CRC) and hepatocellular carcinoma (HCC) ^9, 10^.

**Figure 1.**
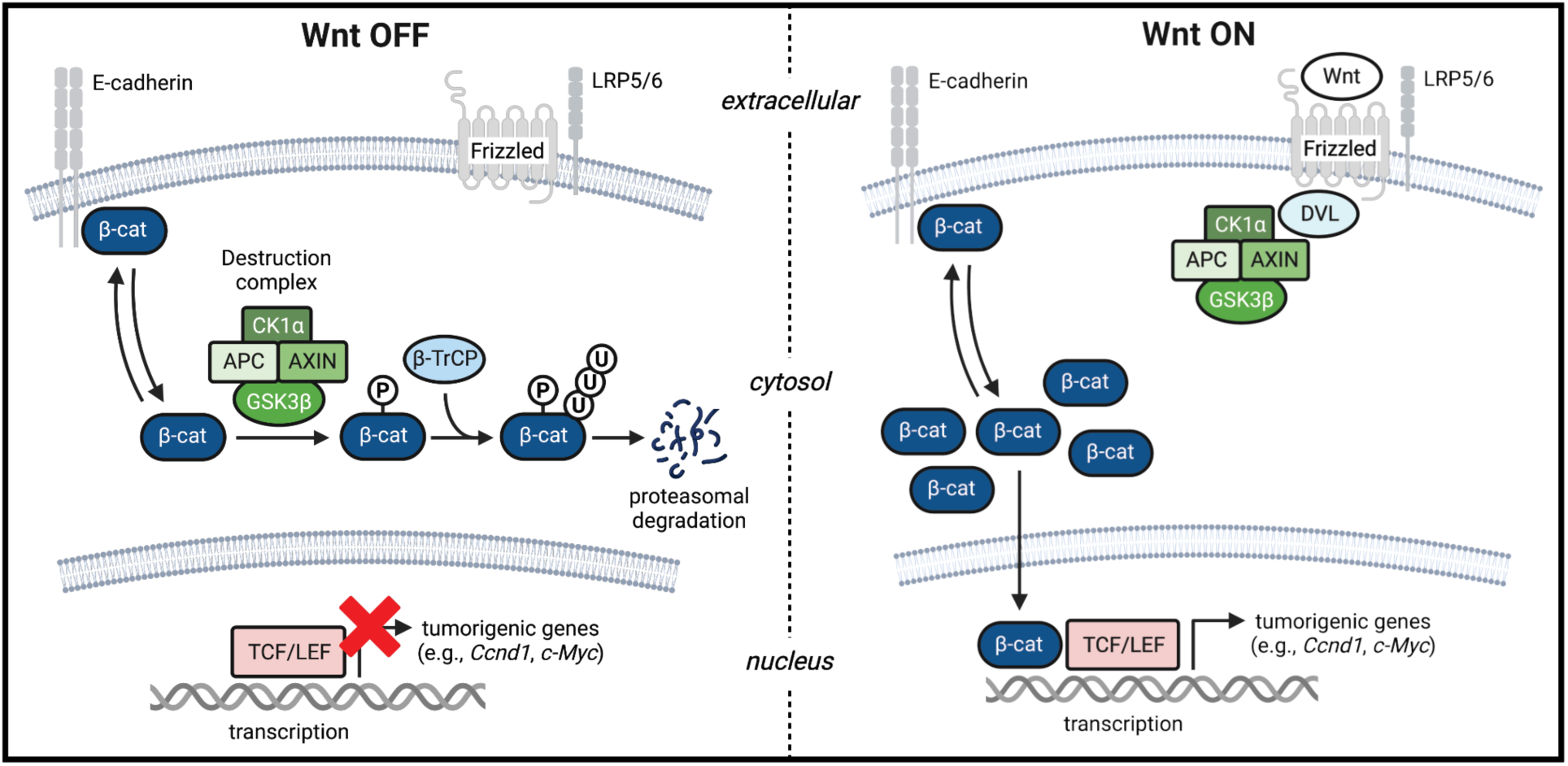
Schematic of canonical Wnt/β-catenin signaling. In the absence of Wnt ligands (left; Wnt “OFF”), the majority of β-catenin is localized at the cytosolic side of the membrane as an integral structural component of E-cadherin-based cell-cell junctions. Free β-catenin in the cytosol is kept at a low level by the activity of the multiprotein destruction complex, which mediates phosphorylation of β-catenin. Phosphorylated β-catenin is ubiquitinated by the E3 ubiquitin ligase β-TrCP and subsequently degraded by the 26S proteasome. In the presence of Wnt ligands (right; Wnt “ON), which interact with a receptor complex consisting of Frizzled protein (FZD) and lipoprotein receptor-related protein 5 (LRP5) or LRP6 on the cell surface, Dishevelled (DVL) and the destruction complex are recruited to the receptor. This recruitment suppresses phosphorylation of β-catenin, which accumulates in the cytosol. Similarly, under pathological conditions, free β-catenin becomes stabilized in the cytosol due to mutations in components of the destruction complex (e.g., truncation mutation in APC such as in DLD1 cells) or in β-catenin directly that prevent it from being phosphorylated and subsequently degraded. Stabilized β-catenin translocates into the nucleus, where it binds T cell factor (TCF) and lymphoid enhancer factor 1 (LEF1) and activates the expression of Wnt target genes in a manner that contributes to the development of various types of cancer.

Given the clearly delineated role of pathogenic β-catenin in tumorigenesis, pharmacological agents designed to prevent abnormal stabilization of β-catenin or facilitate its degradation represent promising approaches that could form the basis of an effective anticancer strategy. Unfortunately, even after decades of preclinical and clinical research, there are currently no approved therapies that target β-catenin directly. Conventional small molecule or monoclonal antibody approaches have met limited success because of β-catenin’s intracellular location, lack of a well-defined, druggable active site, and intrinsically disordered protein regions (IDPRs), which collectively contribute to its classification as an undruggable target ^11^. Beyond direct inhibition, RNA interference (RNAi) approaches such as short interfering RNAs (siRNAs), which silence protein expression at the transcript level, have been developed against β-catenin ^12-14^. However, while siRNA can target any protein-coding mRNA, it has certain limitations. Most critically, siRNA is incapable of distinguishing abnormal β-catenin free in the cytosol and nucleus from normal membrane-associated β-catenin, both of which are expressed from the same gene. This lack of selectivity is potentially problematic because the loss of β-catenin at the membrane decreases cell-cell adhesion, which in turn can promote tumor invasion and metastasis ^15^. Given this and other challenges associated with siRNAs, such as inefficient knockdown of proteins with long half-lives ^16^ and non-specific knockdown of proteins due to partial complementarity with off-target mRNA ^17^, it is evident that a different approach for targeting β-catenin is needed.

One such strategy is proteome editing, a powerful approach for targeted protein modulation that enables post-translational degradation, stabilization, activation, or relocalization of proteins of interest (POIs). By functioning post-translationally, proteome editing has the potential to dissect complicated protein functions at higher resolution than RNAi or gene-editing technologies like CRISPR that operate at the pre-translational level, while also overcoming limitations of these other methods such as irreversibility, lack of temporal control, and off-target effects ^18, 19^. Among the many proteome editing modalities, two of the most advanced are proteolysis targeting chimeras (PROTACs) and molecular glues, which both leverage small molecules to recruit endogenous E3 ubiquitin ligases of the ubiquitin-proteasome pathway (UPP) to POIs for proximity-induced degradation ^20^. However, while a peptide-based PROTAC and molecular glue have been developed for targeted degradation of β-catenin ^21, 22^, they both require very high doses– in the tens of micromolar range–for activity and are incapable of discriminating the cytosolic/nuclear and membrane subpopulations of β-catenin.

An alternative proteome editing approach with the potential to selectively target cytosolic/nuclear β-catenin at lower doses is ubiquibodies (uAbs). Also referred to more recently as affinity-directed protein missiles (AdPROMs) and bioPROTACs, uAbs are chimeric proteins in which an E3 ubiquitin ligase or E3 adaptor is genetically fused with a targeting peptide or protein (warhead) with affinity for a POI. These biologics-based editors redirect otherwise stable POIs to the UPP for proteasomal degradation, as was originally demonstrated with model proteins such as β-galactosidase (β-gal) and green fluorescent protein (GFP) ^23, 24^. Importantly, the modular design of uAbs offers exceptional engineerability, allowing precise customization of both the E3 ligase and the POI-binding warhead ^25^. In the case of E3 ligases, the design space is vast with more than 100 E3s across humans, bacteria and viruses having been functionally incorporated into the uAb architecture ^26-28^, which is in stark contrast to PROTACs that predominantly rely on only two, cereblon and VHL. In the context of targetable proteins, the design space is similarly expansive, taking advantage of an immense collection of available POI-specific scaffolds, such as alpha repeat proteins (αReps), designed ankyrin repeat proteins (DARPins), fibronectin type III (FN3) monobodies, single-chain Fv (scFv) antibodies, VHH nanobodies, and peptides ^23, 24, 26, 29, 30^. Moreover, because uAbs function through protein-protein interactions (PPIs) across extensive contact areas, they can degrade POIs and related proteoforms that have been notoriously difficult to target with conventional small molecule-based modalities ^26, 27, 31-36^. However, POIs that lack pre-existing “off-the-shelf” binding domains, particularly those that are conformationally disordered or devoid of hydrophobic pockets, have been challenging to target using this approach.

To address this challenge, we recently developed several structure-independent protein language models (pLMs) for computationally designing “guide” peptides that enable target engagement by uAb degraders ^29, 37-39^. One such pLM is SaLT&PepPr (Structure-agnostic Language Transformer and Peptide Prioritization) ^29^, a model that leverages fine-tuned ESM-2 pLM embeddings ^40^ to predict the interacting motifs on partner sequences of target POIs and, by integrating with PPI databases, enables isolation of continuous peptide candidates with affinity for an input POI. The resulting SaLT&PepPr-derived peptide warheads were used to construct uAbs, which were experimentally confirmed to bind and degrade their target POIs, including many intrinsically disordered proteins such as regulatory proteins and transcription factors ^29^.

In this study, we leveraged the SaLT&PepPr algorithm to computationally design uAb warheads for driving selective degradation of oncogenic β-catenin in the cytosol and nucleus while preserving membrane-associated β-catenin that is protective and maintains tissue integrity. Specifically, a panel of putative β-catenin-specific uAbs was constructed by fusing SaLT&Pepr-designed guide peptides to the catalytic domain of human C-terminus of Hsp70-interacting protein (CHIP, a.k.a. STUB1), a highly modular human E3 ubiquitin ligase domain that has been used to develop many successful uAbs previously ^23, 29, 30, 38^. Following expression in CRC cells, namely DLD1, that accumulate abnormally high levels of cytosolic β-catenin, we identified several peptide-guided uAb designs that significantly reduced the cytosolic and nuclear pools of β-catenin while sparing the membrane-associated pool. Selective removal of cytosolic and nuclear β-catenin was accompanied by significant inhibition of Wnt/β-catenin signaling activity and impairment of tumor cell survival and proliferation. We also observed selective elimination of cytosolic β-catenin in livers of BALB/C mice that were intravenously injected with lipid nanoparticle (LNP)-encapsulated mRNAs encoding one of the top performing peptide-guided uAbs. Taken together, our findings establish peptide-guided uAbs as a robust proteome editing technology for precisely discriminating between pathogenic and non-pathogenic proteoforms *in vitro* and *in vivo*, with the potential for selectively targeting the primary oncogenic drivers of tumorigenesis.

## RESULTS

### Design of peptide-guided degraders of human β-catenin using SaLT&PepPr

To develop uAbs that selectively eradicate pathogenic β-catenin, we hypothesized that guide peptides based on β-catenin’s known interaction with E-cadherin but with weaker affinity (i.e., >36 nM, which is the measured affinity between the cytosolic domain of E-cadherin (Ecad-CD) and β-catenin ^41^) would preferentially bind cytosolic/nuclear β-catenin while showing minimal competition with the E-cadherin-associated subpopulation at the membrane. To test this hypothesis, we constructed a panel of putative β-catenin-specific uAbs composed of human Ecad-CD-derived guide peptides genetically fused to the catalytic domain of human CHIP that lacked its native substrate-binding domain (CHIPΔTPR) (**Fig. 2a**). Building on our earlier work in which we generated a handful of short, linear β-catenin-specific guide peptides ^29^, here we used the SaLT&PepPr algorithm to generate a larger collection of guide peptides for β-catenin based on its known interaction with Ecad-CD (**Fig. 2b**) ^42^. To this end, the amino acid sequence of Ecad-CD was input to SaLT&PepPr, which first predicted the interaction sites along the Ecad-CD/β-catenin binding interface and then isolated continuous peptide sequences that were scored for their probability of binding to β-catenin. A total of 33 candidate peptides of varying lengths (10−24 amino acids) were generated by this approach (**Supplementary Table 1**). We also leveraged the known interaction between β-catenin and the WD40-repeat domain of the F-box protein β-TrCP (**Fig. 2c**) ^43^ to identify 11 additional peptide candidates (10−20 amino acids) with the potential for β-catenin binding (**Supplementary Table 2**).

**Figure 2.**
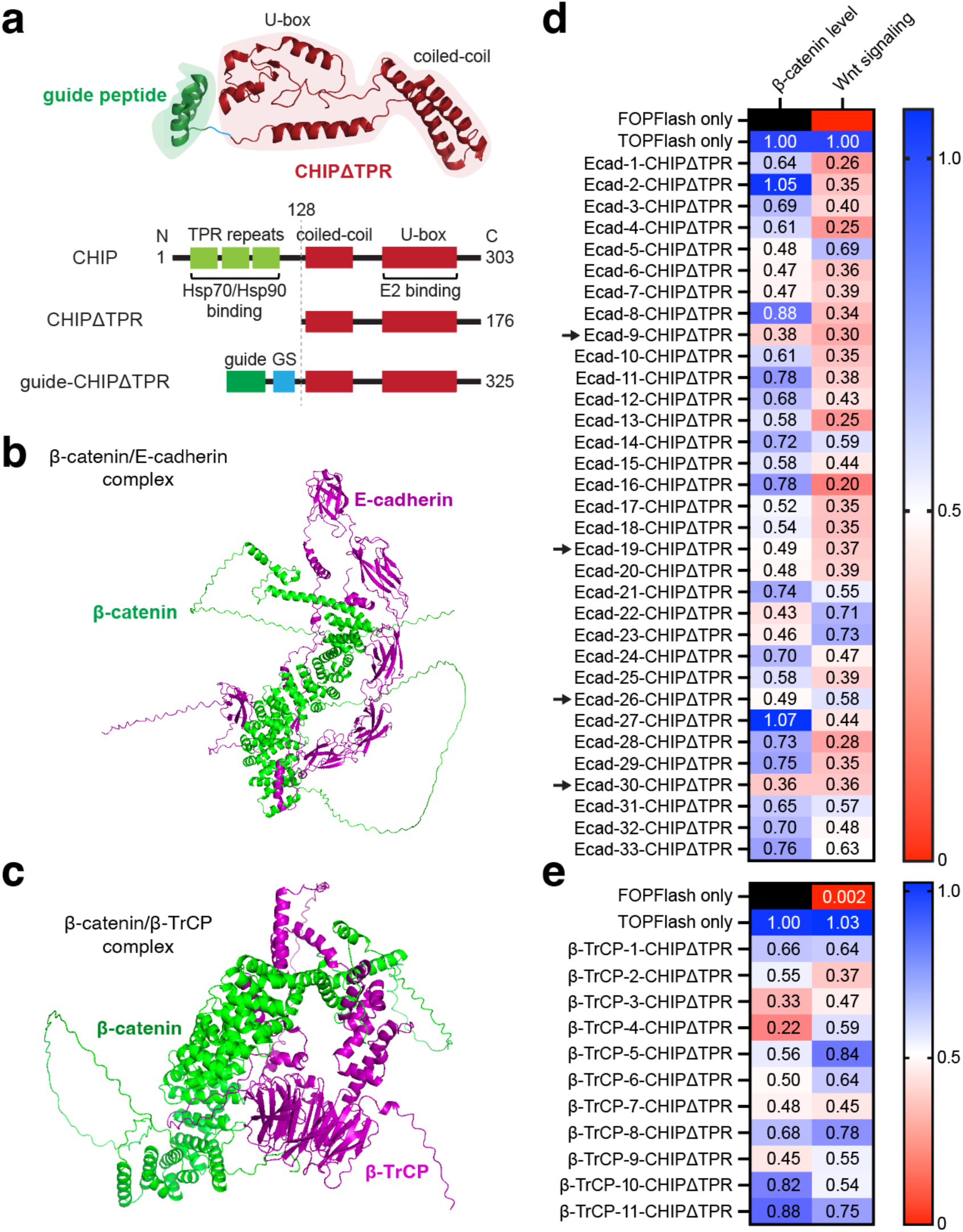
Cytosolic β-catenin knockdown by peptide-guided uAbs. (a) Molecular architecture of peptide-guided uAbs created by removing the N-terminal TPR repeat domain of human CHIP that is responsible for substrate binding and replacing it with a designer guide peptide/protein sequence. In this study, all guides were peptide motifs derived from the SaLT&PepPr algorithm unless otherwise noted. (b,c) Structural models of interaction between β-catenin and either (b) E-cadherin or (c) β-TrCP predicted using AlphaFold3. (d,e) Heatmaps of normalized β-catenin levels determined by densitometry analysis of blots in Supplementary Figures 1a and 2a and normalized β-catenin signaling reported in Supplementary Figures 1b and 2b for uAbs composed of (d) E-cadherin-based and (e) β-TrCP-based peptide guides. Cells receiving only the FOPFlash or TOPFlash plasmid served as negative and positive controls, respectively. Black box = not tested.

### Prioritization of uAbs based on knockdown of β-catenin levels and activity

To test the 44 designs, uAbs were constructed by genetically fusing each guide peptide candidate to CHIPΔTPR in plasmid pcDNA3. We chose to evaluate uAb-mediated β-catenin degradation in DLD1 cells, a CRC cell line in which β-catenin signaling is dysregulated due to loss-of-function mutation in APC, specifically a C-terminal truncation starting at amino acid 1427. This loss of APC canonical function causes aberrant stabilization of β-catenin (**Fig. 1**), which has been associated with a wide variety of human malignancies including CRC ^44^. To determine whether any of the newly designed uAbs could promote the degradation of β-catenin, CRC cells were transiently transfected with uAb-encoding plasmid DNA (pDNA), and cytosolic β-catenin levels were analyzed by immunoblotting. This preliminary screening revealed that all but two of the uAb constructs were capable of lowering steady-state β-catenin levels in the cytosol, with 15 out of 44 promoting a decrease of 50% or more (**Fig. 2d-e** and **Supplementary Figs. 1a, 2a**). In parallel, we assessed the effect of each uAb on β-catenin signaling using TOPFlash, which is a direct and reliable indicator of β-catenin transcriptional activity that involves a TCF/LEF reporter plasmid containing tandemly repeated TCF motifs upstream of the luciferase gene ^32, 45^. Consistent with the immunoblot results, all uAbs reduced luciferase activity to some extent, with several constructs decreasing β-catenin signaling activity by as much as 75−80% (**Fig. 2d-e** and **Supplementary Figs. 1b, 2b**).

Ecad-9-CHIPΔTPR and Ecad-30-CHIPΔTPR were the most effective degraders based on their performance across both immunoblot and TOPFlash screens and thus were down-selected for further analysis. Two additional degraders, Ecad-19-CHIPΔTPR and Ecad-26-CHIPΔTPR, were also down-selected because they showed consistent degradation profiles, and their guide peptides shared a 12-residue core motif (PPYDSLLVFDYE) with the Ecad-9 and Ecad-30 peptides. Importantly, β-catenin knockdown by these peptide-guided degraders was confirmed to be UPP-dependent based on the observation that MG132, an inhibitor of the cytosolic proteasome, completely blocked uAb-mediated degradation of β-catenin (**Supplementary Fig. 3**; shown for Ecad-30-CHIPΔTPR).

### Guide peptides direct binding and ubiquitination of β-catenin *in vitro*

To better understand the functional characteristics of our down-selected, peptide-guided degraders, we evaluated their binding activity and specificity. To this end, each uAb was expressed in *Escherichia coli* strain BL21(DE3) and purified from cell extracts (**Supplementary Fig. 4a**), after which binding was evaluated by enzyme-linked immunosorbent assay (ELISA). As expected, each uAb exhibited strong binding to immobilized β-catenin but not immobilized bovine serum albumin (BSA) (**Supplementary Fig. 4b**), confirming the ability of the computationally designed peptides to direct β-catenin-specific binding. The same purified uAbs were also subjected to biolayer interferometry (BLI) analysis, which revealed the binding affinity (*K*_D_) of each construct to be in the mid-nanomolar range (250-430 nM) (**Supplementary Fig. 4c**), which was ∼10-fold weaker than the reported affinity between β-catenin and Ecad-CD (*K*_D_ = 36 nM) ^41^. For comparison, we generated a panel of uAbs composed of CHIPΔTPR fused to different VHH nanobodies that bind the N-terminal, core, or C-terminal domain of β-catenin with *K*_D_ values ranging from ∼2 nM up to >10 μM ^46^. When expressed in CRC cells, several of the VHH-based uAbs depleted β-catenin with an efficiency that was indistinguishable from the two best peptide-guided degraders, Ecad-9-CHIPΔTPR and Ecad-30-CHIPΔTPR (**Supplementary Fig. 5a-c**). Interestingly, while two of these uAbs were composed of VHHs that have low nanomolar affinity for β-catenin (BC1 and BC2; *K*_D_ ≈ 2−5 nM), two others involved VHHs having order-of-magnitude weaker affinity (BC6 and BC9; *K*_D_ ≈ 5−10 μM) ^46^. Hence, our data indicate that a range of affinities can satisfy the conditions required for strong proximity-induced target degradation of β-catenin and that the guide peptides satisfy our design criteria of binding more weakly to β-catenin than Ecad-CD.

We also investigated the ability of our peptide-guided uAbs to promote the ubiquitination of β-catenin in a reconstituted *in vitro* ubiquitination (IVU) assay. This assay involves mixing purified UPP components (E1, E2, ubiquitin, and ATP) with one of the uAbs as the E3 ubiquitin ligase component and purified β-catenin as the target in a one-pot reaction (**Supplementary Fig. 6a**). It should be noted that β-catenin has 27 potential ubiquitin attachment sites: 26 internal Lys residues as well as its N-terminus. The E2 enzyme UbcH5α was chosen because of its demonstrated ability to function with human CHIP *in vitro* ^23, 30, 47^. When IVU reaction mixtures were probed with an anti-β-catenin antibody, we detected ubiquitinated β-catenin, which appeared in immunoblots as high-molecular-weight (HMW) bands greater than 100 kDa (**Supplementary Fig. 6b**; shown for Ecad-30-CHIPΔTPR). When these samples were immunoprecipitated using magnetic beads coated with a β-catenin-specific VHH nanobody and then immunoblotted with an anti-ubiquitin antibody, HMW ubiquitin species were detected that corresponded to ubiquitinated β-catenin (**Supplementary Fig. 6c**). The intensity of the HMW bands became more pronounced at later incubation times, which was characteristic of the polyubiquitination mediated by CHIP in the presence of native and non-native targets ^23, 30, 47^. In contrast, IVU reactions performed using “guideless” CHIPΔTPR as the E3 showed no detectable ubiquitination of β-catenin, confirming the essentiality of the guide peptide for redirecting the catalytic activity of CHIPΔTPR to the non-native β-catenin target.

To definitively establish the occurrence of β-catenin-linked ubiquitin chains, we profiled the ubiquitination patterns generated by Ecad-30-CHIPΔTPR on β-catenin in IVU reactions. For this analysis, HMW products (∼80−250 kDa) were excised from an SDS-PAGE gel, digested with trypsin, and analyzed by liquid chromatography-tandem mass spectrometry (LC-MS/MS). When a ubiquitinated protein is subjected to tryptic digestion, a C-terminal Gly−Gly dipeptide derived from ubiquitin remains attached to the ubiquitinated Lys residue (**Supplementary Fig. 6d**) ^48^. Therefore, we thoroughly scanned the MS data for the presence of this modification on β-catenin-derived tryptic peptides and identified several ubiquitinated Lys residues in β-catenin (**Supplementary Fig. 6e**), thereby establishing the ability of our peptide-guided uAbs to transfer ubiquitin to multiple Lys residues in the target protein and corroborating the ubiquitin-mediated proteasomal degradation of β-catenin observed above.

### Peptide-guided uAbs selectively degrade cytosolic β-catenin

Having confirmed the specificity and affinity of our peptide-guided degraders, we next investigated their intracellular selectivity for the two subpopulations of β-catenin inside cells, namely the pathogenic cytosolic/nuclear pool and the E-cadherin-bound membrane pool. To this end, we transfected CRC cells with uAb-encoding plasmids or control plasmids and generated subcellular fractions that were subjected to immunoblot analysis. Importantly, cells transfected with peptide-guided uAb degraders exhibited strong, statistically significant knockdown of β-catenin in cytosolic fractions but not in the membrane fractions (**Fig. 3a-b**), indicating clear selectivity of our degraders for the pathogenic signaling pool of β-catenin. In contrast, no measurable degradation was observed in any fractions derived from untreated cells (cells only control) or in control cells transfected with empty pDNA, pDNA encoding the guideless CHIPΔTPR construct, or pDNA encoding a poly-glycine guide peptide fused to CHIPΔTPR (polyG-CHIPΔTPR). For comparison, we also investigated silencing with a small interfering RNA (siRNA) directed against β-catenin (CTNNB1) as per an earlier study ^13^. Interestingly, while both the CTNNB1 siRNA and pDNA-encoded Ecad-30-CHIPΔTPR degrader strongly reduced β-catenin levels in the cytosolic fraction of CRC cells, the siRNA also significantly lowered the membrane pool of β-catenin (**Supplementary Fig. 7**) and thus lacked the selectivity of our peptide-guided uAbs.

**Figure 3.**
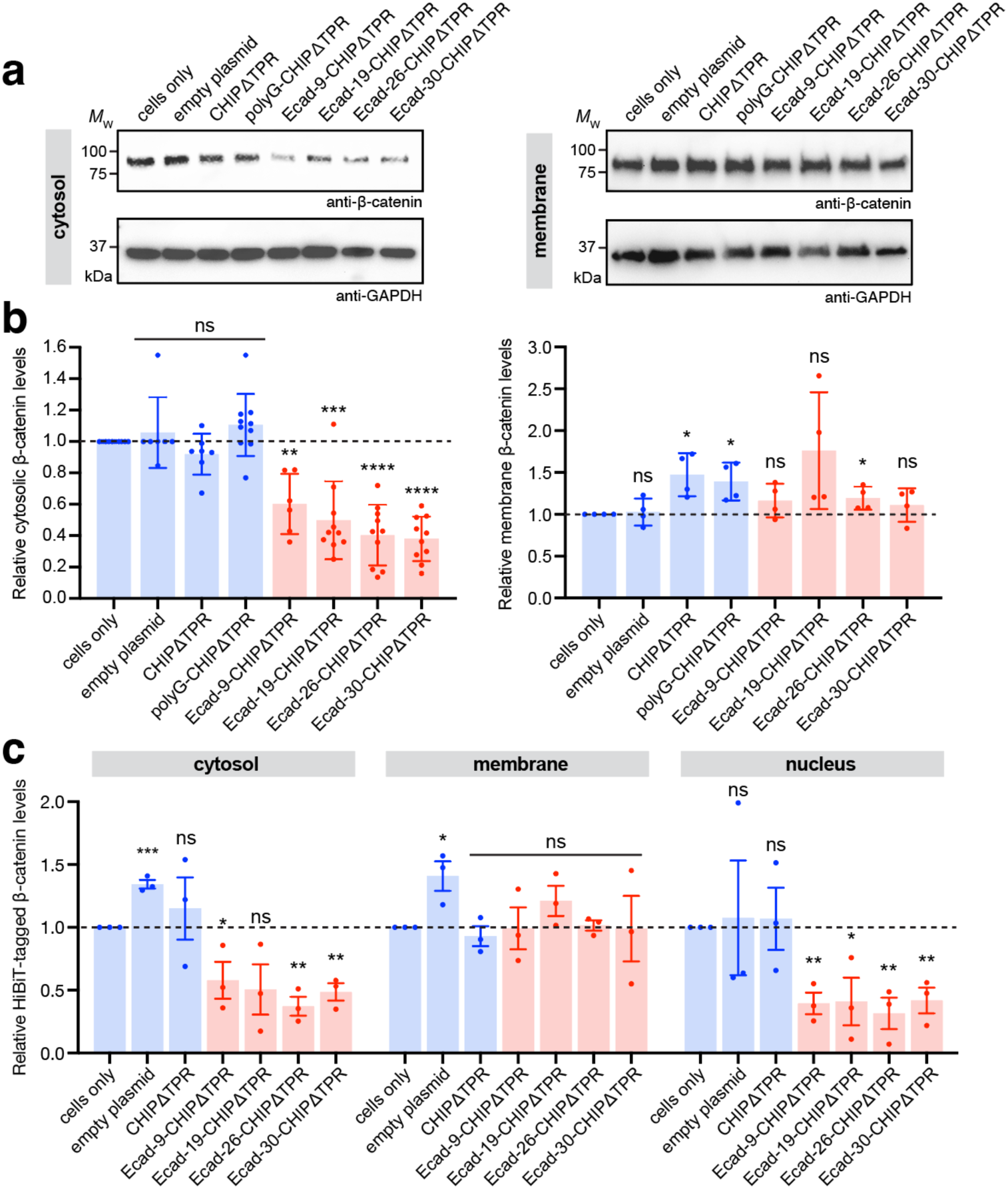
Selective degradation of cytosolic/nuclear but not plasma membrane-associated β-catenin. (a) Immunoblot analysis of cytosolic and membrane β-catenin levels in DLD1 cells transfected with empty plasmid pcDNA3 or pcDNA3 encoding one of the peptide-guided uAb degraders, CHIPΔTPR, or polyG-CHIPΔTPR. Cells were harvested 48 h post-transfection, after which cytoplasmic and membrane fractions were prepared from cell extracts and subjected to immunoblotting with anti-β-catenin antibody (top) and anti-GAPDH antibody (bottom), the latter serving as a loading control for both cytosolic and membrane fractions. Lanes were normalized by total protein content and molecular weight (MW) markers are indicated at left. Blots are representative of at least three biological replicates. (b) Quantification of cytosolic and membrane β-catenin levels by densitometry analysis of immunoblots in panel (a). Band intensity was determined using ImageJ software with all β-catenin band intensities normalized to corresponding GAPDH band intensities. Relative β-catenin levels were then calculated by normalizing all values to cells only control. Data are mean of at least three biological replicates (*n* = 3−6) ± SD. (c) HiBiT-based quantification of β-catenin levels in cytosolic, membrane, and nuclear fractions derived from DLD1 cells transfected with empty plasmid pcDNA3 or pcDNA3 encoding one of the peptide-guided uAbs or CHIPΔTPR. Relative HiBiT-tagged β-catenin levels were calculated by normalizing all values to cells only control. Data are mean of three biological replicates (*n* = 3). Statistical significance was determined by unpaired two-tailed Student’s *t-*test. Calculated *p* values are represented as follows: *, *p* < 0.05; **, *p* < 0.01; ***, *p* < 0.001; ****, *p* < 0.0001; ns, not significant.

To probe the uAb selectivity more quantitatively, we took advantage of a CRISPR/Cas9 edited DLD1 cell line in which a sequence encoding the 11 amino acid HiBiT tag was knocked-in at the endogenous *CTNNB1* loci, resulting in C-terminal tagging of the β-catenin protein. Following transfection of the HiBiT reporter cells with the peptide-guided uAbs, we analyzed subcellular fractions and observed strong knockdown (∼50−60%) of cytosolic β-catenin but no measurable change in the membrane β-catenin levels (**Fig. 3c**), consistent with the immunoblotting results. It should be noted that cytosolic and membrane β-catenin levels remained elevated in the cells only control or cells transfected with empty pDNA or pDNA encoding guideless CHIPΔTPR. We further tested nuclear fractions derived from the same cells and observed that all degraders promoted significant reduction of β-catenin levels in the nucleus whereas none of the controls showed any evidence of β-catenin knockdown (**Fig. 3c**), mirroring the cytosolic β-catenin profiles. Collectively, these findings indicate that our peptide-guided uAbs preferentially degraded the soluble cytosolic/nuclear subpopulation of β-catenin while sparing the membrane-bound subpopulation of β-catenin, which is tightly associated with endogenous E-cadherin at the cell membrane.

### Functional impact of uAb-mediated degradation of cytosolic/nuclear β-catenin

Given the strong depletion of the cytosolic/nuclear signaling pool of β-catenin, we next investigated the effect of our peptide-guided degraders as well as control constructs on the transcriptional activity of β-catenin using the TOPFlash reporter in CRC cells. In agreement with the immunoblot and HiBiT results, all four uAbs significantly inhibited β-catenin signaling as evidenced by a strong reduction in luciferase activity, which was on par with that observed in CRC cells transfected with the CTNNB1 siRNA (**Fig. 4a**). In contrast, the cells only control or cells transfected with control pDNA showed no reduction in β*-*catenin signaling. Next, we determined if uAb-mediated down-regulation of β-catenin signaling corresponded to decreased expression of known β-catenin target genes, namely *Axin2* ^49^, *Cyp1a2* ^50^ and *c-Myc* ^6^. Consistent with the TOPFlash results, real-time quantitative PCR (qPCR) analysis revealed a significant decrease in expression of the β-catenin target genes in CRC cells transfected with pDNA encoding the Ecad-30-CHIPΔTPR construct but not in CRC cells transfected with control pDNA (**Fig. 4b**). Taken together, these results confirm the ability of peptide-guided uAb degraders to induce a pronounced loss of β-catenin function in tumor cells.

**Figure 4.**
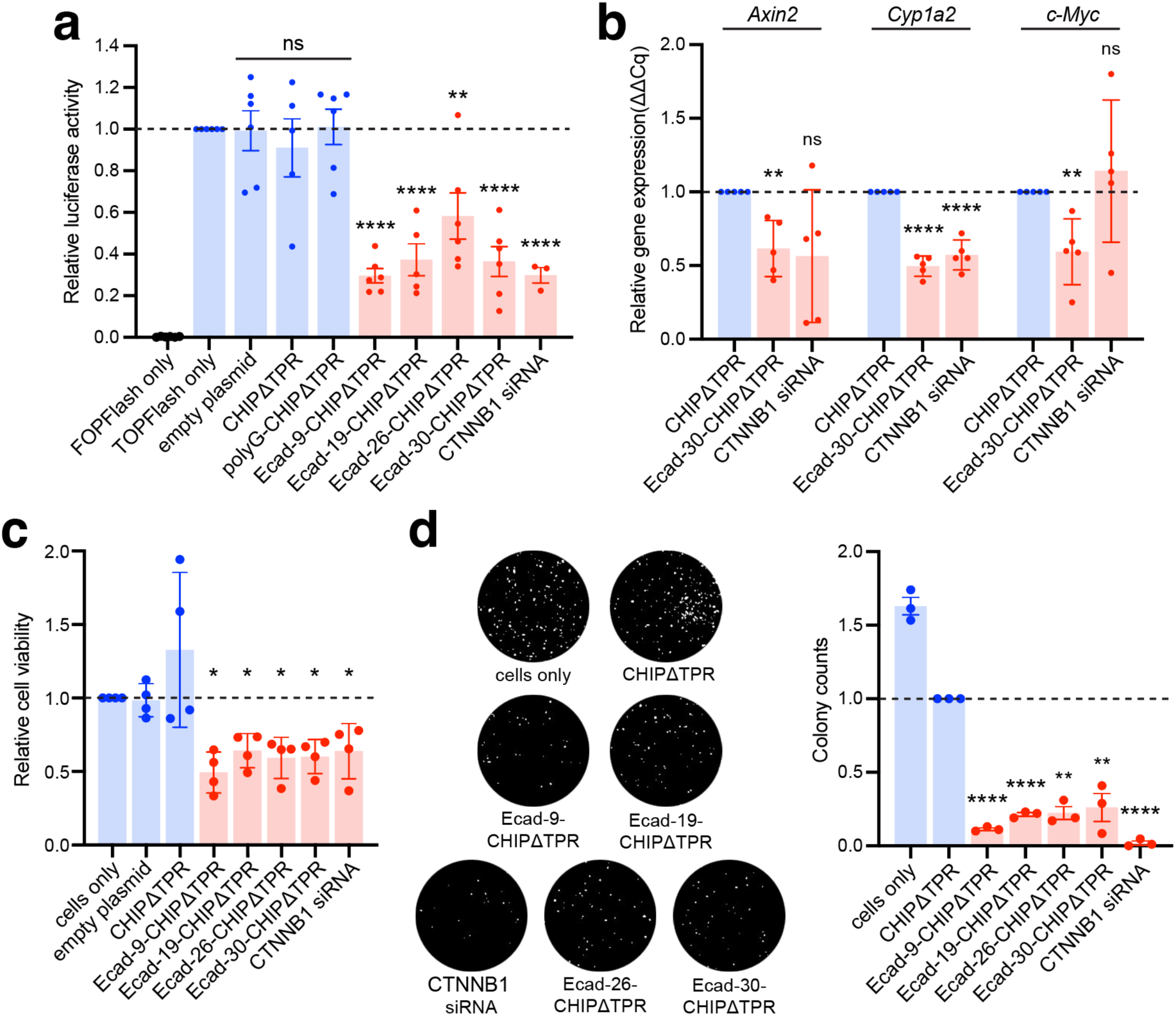
Functional impact of cytosolic/nuclear β-catenin knockdown by peptide-guided uAbs. (a) β-catenin signaling activity in DLD1 cells co-transfected with TOPFlash reporter plasmid along with empty plasmid pcDNA3, pcDNA3 encoding one of the peptide-guided uAb degraders, CHIPΔTPR, or polyG-CHIPΔTPR, or CTNNB1 siRNA. Cells receiving only the FOPFlash or TOPFlash plasmid served as additional negative and positive controls, respectively. Luciferase signals in each sample were normalized to those measured in control cells receiving no degrader plasmid. Data are mean of three or more biological replicates (*n* = 3−6) ± SD. (b) qPCR analysis of known β-catenin target genes, *Axin2*, *Cyp1a2* and *c-Myc*, in DLD1 cells transfected with pcDNA3 encoding CHIPΔTPR or Ecad-30-CHIPΔTPR or transfected with CTNNB1 siRNA. Relative gene expression was normalized by the ΔΔCq method with *Gapdh* as the reference gene; these values were subsequently normalized to signal for CHIPΔTPR. Data are mean of biological replicates (*n* = 3) ± SD. (c) Viability of DLD1 cells transfected with empty plasmid pcDNA3, pcDNA3 encoding CHIPΔTPR, or one of the peptide-guided uAb degraders, or CTNNB1 siRNA. Cells were harvested 48 h post-transfection, after which viability was quantified by MTS assay. Data are mean of biological replicates (*n* = 4) ± SD. (d) Cell proliferation assay for cells in (c). Colony-forming ability was assessed by diluting cells, plating at a low density, and allowing to grow for 5 days. Plates were photographed (left panel) and the number of crystal violet-stained colonies was counted using the ImageJ software (right panel). Data are mean of biological replicates (*n* = 3) ± SD. Statistical significance in all panels was determined by unpaired two-tailed Student’s *t*-test. Calculated *p* values are represented as follows: *, *p* < 0.05; **, *p* < 0.01; ***, *p* < 0.001; ****, *p* < 0.0001; ns, not significant.

To determine the biological impact of uAb-mediated β-catenin reduction, tumor cells were examined for viability by MTS assay and proliferation by colony formation assay. In terms of viability, a significant effect on survival was apparent in CRC cells at 24 and 48 h after transfection with pDNA encoding the Ecad-30-CHIPΔTPR construct but not control pDNA (**Fig. 4c** and **Supplementary Fig. 8**). The magnitude of this uAb-mediated decrease in cell viability (∼35-50%) was on par with that measured for tumor cells transfected with CTNNB1 siRNA (∼35%) and in close agreement with a previous report ^13^. Next, we examined the effect of β-catenin loss on tumor cell proliferation. Following transfection with pDNA encoding the uAbs or guideless CHIPΔTPR construct, cells were diluted, plated at a low density, and cultured in plates for 5 days. A dramatic decrease in crystal violet-stained colonies was evident only in the uAb-transfected CRC cells, as shown in representative cultures, with uAbs inducing statistically significant decreases in the number of colonies relative to control constructs (**Fig. 4d-e**). It should be noted that the marked reduction in colony forming capability induced by the uAbs was comparable to that induced by transfection with the CTNNB1 siRNA. Collectively, these data indicate that the robust functional knockout of β-catenin triggered by our peptide-guided uAbs has clear functional consequences on tumor cell viability and proliferation.

### Lipid-mediated intracellular delivery of mRNAs encoding peptide-guided uAbs

While uAb degraders offer a promising proteome editing strategy, efficient cellular delivery represents a major obstacle that limits their therapeutic potential. To address this challenge, we investigated the delivery of uAb-encoded mRNAs using LNPs, which have emerged as reliable, therapeutically relevant vectors for targeted delivery, cellular uptake, and cytosolic release of RNA payloads. Importantly, LNP components are widely regarded as safe, and FDA approval has been granted for LNP formulations that encapsulate mRNA and siRNA ^51-53^.

As an initial proof-of-concept of the strategy, synthetic mRNAs encoding Ecad-30-CHIPΔTPR mRNA, guideless CHIPΔTPR, and CHIPΔTPR fused with a non-specific scramble peptide (scr-CHIPΔTPR) were produced by *in vitro* translation (IVT) and subsequently formulated with LNPs in which the ionizable lipid was a novel trialkyl ionizable lipid called Lipid 10 ^54^. We chose Lipid 10 because it is a well-tolerated and potent ionizable lipid for siRNA and mRNA delivery in rodents and nonhuman primates (NHPs) ^54^. Moreover, unlike some other recently disclosed ionizable lipids (e.g., SM-102 and ALC-0315 used in COVID vaccines), Lipid 10 was designed for intravenous (i.v.) delivery and found to perform optimally for targeting hepatocytes ^54^.

Because this LNP composition was expected to drive biodistribution to the liver following i.v. administration, we first evaluated our uAb-mRNA-LNP formulations using the human HCC cell line, Hep3B, which is known for its extensive expression of liver-specific proteins and abnormally stabilized β-catenin levels due to mutation of *AXIN1*. Transfection of Hep3B cells with Ecad-30-CHIPΔTPR-mRNA-LNP resulted in strong knockdown of cytosolic β-catenin whereas treatment with PBS or control LNP formulations resulted in no detectable knockdown (**Supplementary Fig. 9a-b**). It is worth noting that this level of β-catenin knockdown in the cytosol was on par with that observed in Hep3B cells transfected with pDNA encoding the same degrader (**Supplementary Fig. 9c-d**). We also evaluated the effect of LNP-delivered uAb mRNA on β-catenin signaling. Specifically, Hep3B cells transfected with the TOPFlash reporter were treated with LNPs formulated with increasing concentrations of mRNA encoding Ecad-30-CHIPΔTPR or control constructs. Consistent with the significant knockdown of cytosolic β-catenin, delivery of Ecad-30-CHIPΔTPR-mRNA-LNP resulted in robust and dose-dependent inhibition of β*-*catenin signaling, with a half-maximal inhibition (IC_50_) of 2.9 nM at 48 h post-treatment and an overall reduction in luciferase activity that rivaled that measured in Hep3B cells transfected with pDNA encoding Ecad-30-CHIPΔTPR (**Supplementary Fig. 10a-c**). As expected, treatment with PBS or LNP-encapsulated control constructs did not elicit any changes in luciferase activity.

### LNP-delivered uAb mRNA silences cytosolic β*-*catenin in mice

Given the ability of Ecad-30-CHIPΔTPR-mRNA-LNP to promote knockdown of cytosolic β-catenin *in vitro*, we next investigated whether the same formulations could promote knockdown of β-catenin *in vivo* following systemic administration. To this end, groups of wild-type BALB/c mice were injected intravenously in the lateral tail vein with a single injection of PBS or a 1.0 mg/kg dose of the same uAb-mRNA-LNP formulations described above. At 24 h after the i.v. injections, we analyzed tissue-specific β*-*catenin silencing by homogenizing isolated liver tissue and performing subcellular fractionations to generate cytosolic and membrane fractions. As expected, LNP-delivered mRNA encoding CHIPΔTPR or scr-CHIPΔTPR control constructs showed no measurable changes in cytosolic or membrane β*-*catenin levels relative to the levels measured in mice receiving PBS (**Fig. 5a-b**). Meanwhile, LNP-mediated delivery of Ecad-30-CHIPΔTPR mRNA resulted in statistically significant elimination of cytosolic β*-*catenin but not membrane-associated β*-*catenin (**Fig. 5a-b**). These results mirrored the strong and selective knockdown of cytosolic but not membrane-associated β*-*catenin observed in cultured hepatoma cells, thereby confirming that the selectivity of our peptide-guided degrader was maintained following LNP-mediated uAb mRNA delivery *in vivo.* The duration of this effect was evaluated in a short-term study by examining time course changes in liver-specific β*-*catenin levels by immunoblotting analysis. Specifically, liver samples were harvested on days 1–7 after a single 1-mg/kg dose of Ecad-30-CHIPΔTPR-mRNA-LNP or CHIPΔTPR-mRNA-LNP. Notably, in mice receiving the Ecad-30-CHIPΔTPR-mRNA-LNP formulation, cytosolic β*-* catenin remained selectively decreased on days 1–5 post-injection but returned to steady-state levels by day 7, while membrane β-catenin levels were unchanged over this same interval (**Supplementary Fig. 11**). In contrast, mice receiving the CHIPΔTPR-mRNA-LNP control formulation exhibited no measurable changes in cytosolic or membrane β*-*catenin levels. Collectively, these results lay the foundation for the potential clinical translation of peptide-guided β*-*catenin degraders in pre-clinical and clinical models of Wnt-driven CRC and HCC.

**Figure 5.**
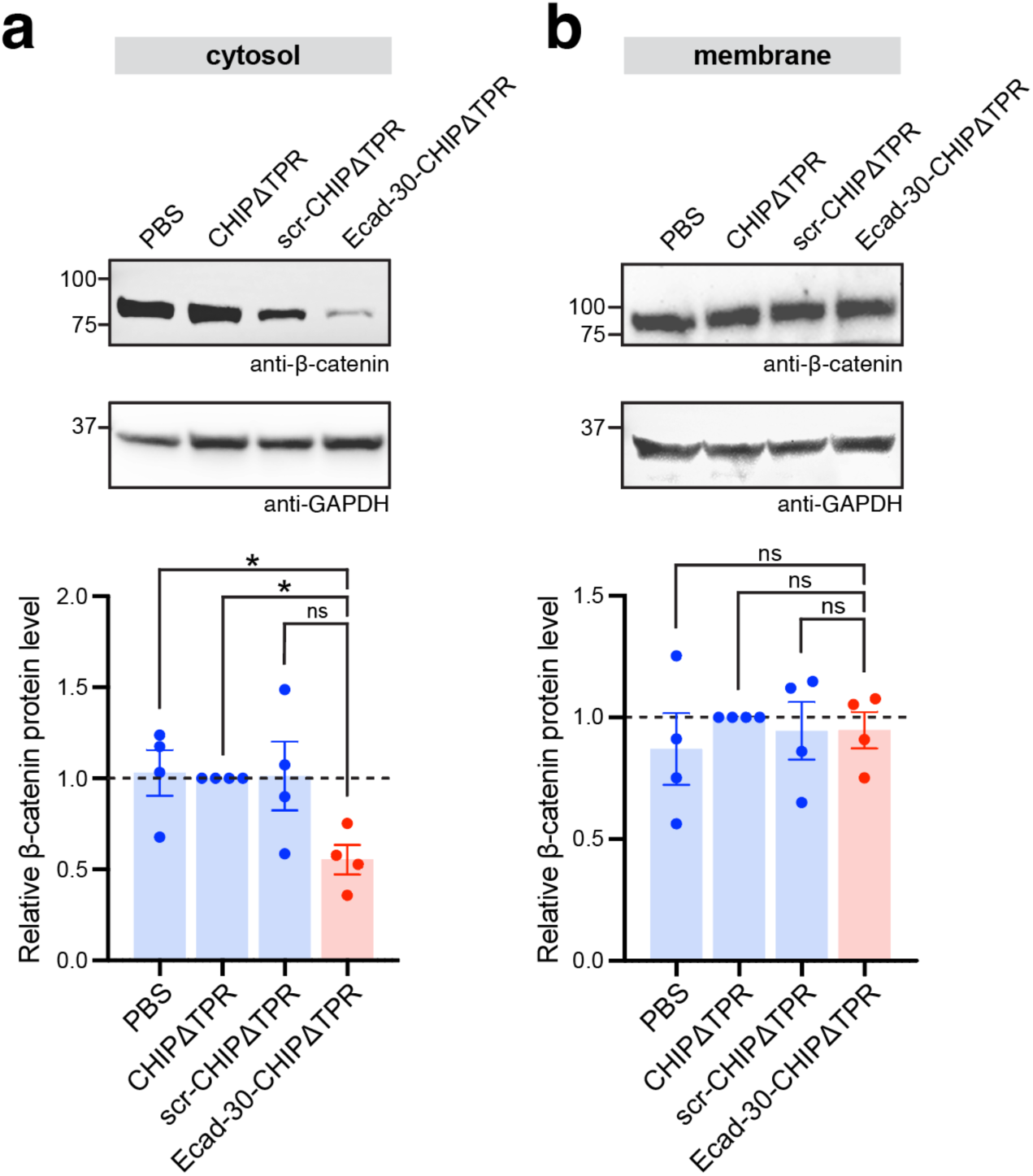
Silencing of cytosolic β*-*catenin following LNP-mediated delivery of uAb mRNA in mice. (a) Immunoblot analysis of β-catenin in cytosolic (left) and membrane (right) fractions derived from homogenized livers of wild-type BALB/c mice (*n* = 4 mice per group). Mice were injected intravenously with a single 1.0 mg/kg dose of the following: PBS, CHIPΔTPR-mRNA-LNP, scr-CHIPΔTPR-mRNA-LNP, or Ecad-30-CHIPΔTPR mRNA-LNP. Blots were probed with anti-β-catenin antibody (top) and anti-GAPDH antibody (bottom), the latter serving as a loading control for both cytosolic and membrane fractions. Lanes were normalized to the tissue weight and total protein for each liver. Molecular weight (MW) markers are indicated at left. Blots are representative of two technical replicates per mouse. (b) Quantification of cytosolic and membrane β-catenin levels by densitometry analysis of immunoblots in panel (a). Band intensity was determined using ImageJ software with all β-catenin band intensities normalized to corresponding GAPDH band intensities. Relative β-catenin levels were then calculated by normalizing values to CHIPΔTPR control. Data are mean of biological replicates ± SD, where each data point is the average of two technical replicates. Statistical significance was determined by unpaired two-tailed Student’s *t*-test. Calculated *p* values are represented as follows: *, *p* < 0.05; **, *p* < 0.01; ns, not significant.

## DISCUSSION

Here, we describe a method for rapidly designing uAb degraders in a CRISPR-analogous manner whereby short guide peptides were identified using a structure-agnostic pLM, SaLT&PepPr, and used to redirect the human E3 ubiquitin ligase CHIP to pathogenic β-catenin and accelerate its removal via proteasomal degradation. Importantly, our peptide-guided uAb degraders were shown to selectively eradicate abnormally accumulated β-catenin in the cytosol and nucleus of CRC cells while preserving normal β-catenin at the membrane. In contrast, a β-catenin-targeting siRNA was unable to distinguish between these subpopulations, as they are expressed from the same gene. This distinction arises because, unlike RNAi and CRISPR-Cas methods, uAb-mediated proteome editing operates at the post-translational level. Consequently, uAbs have the potential to dissect complicated protein functions with higher resolution than their gene silencing counterparts. For example, by exploiting the customizable affinity and specificity of the warhead, uAbs have been engineered that selectively deplete particular protein states (*e.g*., active/inactive conformation, mutant/wildtype, post-translationally modified/unmodified, etc.) as well as localizations, as we showed here and has been reported previously ^26, 27, 30-34, 36, 55, 56^.

In the present study, we selectively targeted the pathogenic subpopulation of β-catenin by designing guide peptides based on β-catenin’s known interaction with E-cadherin but with weaker affinity than the native interaction. The resulting uAbs showed minimal competition with the E-cadherin-associated subpopulation at the membrane. Interestingly, these uAbs exhibited a mid-nanomolar affinity (250-430 nM) for β-catenin, which is measurably weaker than the low nanomolar affinity of warheads that are commonly deployed in uAb studies. In fact, our peptide-guided uAbs performed as well or better than uAbs constructed with low nanomolar affinity (∼2-5 nM) VHH domains specific for β-catenin, consistent with a previous study in which a DARPin-CHIPΔTPR chimera with low micromolar affinity for ERK2 was still capable of achieving target degradation ^30^. Collectively, these results further emphasize that warheads with high affinity may not be optimal for constructing an efficacious uAb ^34^ and that other parameters in addition to binding kinetics, such as cooperativity, dynamics, epitope differences, and proximity considerations, are crucial determinants of ternary complex formation and ubiquitination efficiency ^57^. In the future, it will be important to more carefully map the relationship between affinity and efficacy, while also keeping in mind that naturally occurring E3 ligases often exhibit relatively modest affinities for their targets. For example, the measured affinity between CHIP and its native substrates Hsp70, Hsp90, and Hsc70 is in the low micromolar range (*K*_D_ = 0.3−2.3 μM) ^58^.

A key design feature that was leveraged here is the modularity of human CHIP, which derives from its conformational flexibility ^47^ and broad substrate specificity ^59^. This modularity allows CHIP to accommodate substantial rewiring of its target-binding domain without compromising its ubiquitin transfer activity, making it widely applicable as a versatile degrader of diverse POIs in both transiently and stably transfected cell lines, as evidenced by the many successful uAbs that have enlisted this E3 ^23, 27, 29, 30, 36, 38, 60-62^. Indeed, simple swapping of the warhead has proven to be an effective means for on-demand construction of functional uAbs targeting new POIs ^25, 57, 63^. However, while the exchange of uAb warheads is relatively straightforward, the discovery of entirely new ones (i.e., beyond pre-existing “off-the-shelf” binders) has remained a significant bottleneck. This process has historically relied upon purely experimental methods, such as animal immunization or screening using display technologies (e.g., phage, yeast, etc.), which are time-consuming, labor-intensive, and often result in low hit rates. Fortunately, recent breakthroughs in generative methods have revolutionized *de novo* target binder design and quickened the pace of discovery. State-of-the-art pLMs, such as ESM-2, now offer unprecedented potential to create binders for virtually any POI, requiring little to no structural information ^64^. Consequently, structure-agnostic approaches like SaLT&PepPr ^29^, PepPrCLIP ^37^, and PepMLM ^39^ have proven particularly effective at designing binders that target disordered proteins, including difficult-to-drug transcription factors like β-catenin. Moreover, pLM-driven binder discovery has greatly accelerated the pace at which new uAbs can be generated, as demonstrated here and in other recent studies ^29, 37-39^. With SaLT&PepPr, we rapidly constructed a panel of 44 bespoke uAbs that were evaluated for their ability to degrade pathogenic β-catenin and inhibit its signaling activity, with design-build-test cycles of roughly one week. This development speed is in stark contrast to small molecule-based degraders (e.g., molecular glues, PROTACs), which suffer from the lack of available POI- and E3-specific ligands and typically require extremely long and difficult campaigns to discover ligands for new targets ^25^. It is also worth noting that, although we focused on four of the best uAb degraders for in-depth characterization, our preliminary screen identified 29 out of 44 uAb designs that promoted >30% reduction in both β-catenin levels and Wnt/β-catenin signaling activity, reflecting a 66% success rate.

While peptide-guided uAbs hold great potential for knockdown of biomedically important targets, their clinical application is limited by the fact that most protein biologics are incapable of spontaneously entering mammalian cells ^65^. Consequently, methods for efficient *in vivo* delivery are needed for uAbs to reach their full clinical potential. However, most studies that have evaluated uAb efficacy *in vivo* have relied on delivery methods that are unsuitable for clinical translation, such as stable transfection/transduction of tumor-cell lines with uAb genes prior to implantation in mice ^31-34, 55, 62^ or injection of recombinant adenovirus delivery vehicles via non-clinically relevant routes of administration (e.g., intra-tumoral, intra-amniotic) ^61, 66-68^. With an eye towards more clinically relevant, non-viral delivery strategies, LNP vehicles have been used to encapsulate anionically-modified uAb proteins, leading to efficient intracellular delivery and target knockdown in cultured cells ^69^; however, whether this method is effective *in vivo* was not tested. In related work, our group demonstrated the use of cationic polypeptide-based nanoplexes to functionally deliver encapsulated uAb-encoding mRNAs to mice ^26^; however, this formulation has yet to be clinically validated.

Here, we pursued a well-established path for enabling exogenous proteins to access intracellular targets by delivering their encoding mRNA via LNP carriers, which are one of the most advanced non-viral delivery systems and are approved for therapeutic use in humans ^51-53^. We found that uAb-mRNA-LNP formulations enabled highly selective removal of pathogenic β-catenin *in vitro*, resulting in potent suppression of Wnt/β-catenin signaling with doses of less than 10 nM. When the same formulations were administered in mice, we observed strong knockdown of cytosolic β-catenin that persisted for 5 days following i.v. injection of a 1.0 mg/kg dose, consistent with the duration of protein expression reported in other single-dose experiments where mRNA-LNP formulations were i.v. injected at comparable mRNA concentrations ^70, 71^. To further extend the duration of uAb-mediated knockdown, several strategies could be envisioned. Besides improvements to mRNA payloads and LNP vehicles themselves, which have become areas of intense research ^72^, uAb-centric innovations will also be crucial for *in vivo* therapeutic applications. For example, nearly all E3s are subject to autoubiquitination, which is an essential part of their natural turnover but can also lead to self-destruction of uAbs following their cytosolic delivery ^31, 34, 69^. Thus, lysine-replacement strategies that render uAbs resistant to self-degradation without compromising specificity and ubiquitination efficiency ^23, 34^ (US Patent Application No. 18/845,621) will need to be integrated with *in vivo* delivery efforts in the future.

As uAb degraders and LNP delivery vehicles are further refined and optimized, we anticipate that uAbs will become an increasingly attractive modality for targeting intracellular proteins, especially given the remarkable pace with which novel warheads are being discovered using pML-driven algorithms ^29, 37-39^. Our demonstration of LNP-mediated delivery of these peptide-guided degraders into cells, both *in vitro* and *in vivo*, serves as an important proof-of-concept for translating the uAb technology platform and sets the stage for uAb-mediated treatment of challenging diseases in the future.

## MATERIALS AND METHODS

### Computational peptide design

Binding peptides designed in this study were generated by inputting the Ecad-CD or β-TrCP interacting partner sequences (**Supplementary Table 3**) into the SaLT&PepPr algorithm ^29^ (https://huggingface.co/ubiquitx/saltnpeppr). All binder sequences can be found in Supplementary Tables 1 and 2.

### Plasmids

For construction of all peptide-guided uAb plasmids, a pcDNA3 vector containing an Esp3I restriction site immediately upstream of DNA encoding a flexible GSGSG linker followed by the CHIPΔTPR gene was used ^29^. Oligonucleotides encoding the candidate guide peptides were annealed and subsequently ligated into the Esp3I-digested backbone using T4 DNA ligase (NEB). Assembled constructs were used to transform *E. coli* cells (DH5α) and plated onto Luria-Bertani (LB)-agar supplemented with the appropriate antibiotic. For protein purification, genes encoding each uAb construct were PCR-amplified with primers that introduced a C-terminal 6x-His tag and subsequently cloned into plasmid pET28a between the XbaI and EcoRI restriction sites. All plasmids were isolated using a QIAprep Spin Miniprep Kit (Qiagen) and confirmed by DNA sequencing at the Genomics Facility of the Cornell Biotechnology Resource Center (BRC).

### Cell culture and transfection

The human CRC cell line, DLD1, was purchased from ATCC (cat # CCL-221). DLD1 cells were cultured in RPMI1640 media (ThermoFisher) supplemented with 100 units/mL penicillin, 100 mg/mL streptomycin, and 10% FBS (Corning) at 37 °C with 5% CO_2_. Cells were seeded in a 6-well plate one day before transfection. Plasmids encoding the peptide-guided uAb constructs were prepared using the PureYield miniprep kit (Promega) to eliminate endotoxins. An appropriate amount of plasmid DNA was mixed with jetPRIME buffer by vortexing for 10 s, after which jetPRIME reagent (VWR) was added to the mixture and incubated for 10 min at room temperature. The mixture was then added to the cells. After 4 h of incubation at 37 °C, with 5% CO_2_, fresh media was replenished. Cells were subsequently cultured under these conditions for an additional 48 h before being harvested for analysis. For siRNA transfection, a final concentration of 100 nM CTNNB1 siRNA (Cell Signaling, cat # 6225) was transfected in each well of a 96-well plate (100 μl per well) using jetPRIME reagent.

### Immunoblot analysis

On the day of harvest, cells were detached by addition of 0.05% trypsin-EDTA (ThermoFisher) and cell pellets were washed twice with ice-cold 1x PBS. Cells were then lysed, and subcellular fractions were isolated from lysates using a Subcellular Protein Fractionation Kit (ThermoFisher) per the manufacturer’s instructions. Specifically, ice-cold cytosolic extraction buffer was added to the cell pellet, the mixture was placed at 4 °C for 10 min with gentle shaking followed by centrifugation at 500 × g for 10 min at 4 °C. The supernatant was collected immediately to a pre-chilled PCR tube and placed on ice followed by immunoblotting or stored at –20 °C for future usage. The pellet was then mixed with ice-cold membrane extraction buffer and the mixture was incubated at 4 °C for 10 min followed by centrifugation at 3000 × g for 5 min. The supernatant was immediately transferred to a pre-chilled tube. Protein concentration was quantified using the Pierce BCA Protein Assay Kit (ThermoFisher). An equivalent amount of total protein was loaded into Precise Tris-HEPES 4–20% sodium dodecyl sulfate (SDS)-polyacrylamide gels (ThermoFisher) and separated by electrophoresis. Immunoblotting was performed according to standard protocols. Briefly, proteins were transferred to poly(vinylidene fluoride) (PVDF) membranes (Millipore), blocked with 5% (w/v) nonfat dry milk (Carnation) in 1x tris-buffered saline (TBS) with 0.05% (v/v) Tween 20 (TBST) at room temperature for 1 h, washed three times with TBST for 10 min, and probed with rabbit anti-β-catenin (Cell Signaling, cat # 8480S; diluted 1:1000); mouse anti-GAPDH (Calbiochem, cat # CB1001; diluted 1:5000); or rabbit anti-β-tubulin (Cell Signaling, cat # 2146; diluted 1:1000). The blots were washed again three times with TBST for 5 min each and then probed with a secondary antibody, either donkey anti-rabbit-HRP (Abcam, cat # ab7083; diluted 1:2500) or goat anti-mouse-HRP (Abcam, cat # ab97023; diluted 1:4000) for 1 h at room temperature. Blots were detected by chemiluminescence using a ChemiDoc MP imager (Bio-Rad). Densitometry analysis of protein bands in immunoblots was performed using ImageJ software ^73^ as described at https://imagej.net. Briefly, bands in each lane were grouped as a row or a horizontal “lane” and quantified using the gel analysis function in ImageJ. Intensity data for the uAb bands was normalized to band intensity for CHIPΔTPR control unless stated otherwise.

### TOPFlash assay

A total of 1 × 10^4^ DLD1 cells were seeded on a white-bottom 96-well plate 24 h prior to transfection. On the day of transfection, each well received the following plasmids: M50 Super 8x TOPFlash plasmid (Addgene plasmid # 12456) or M51 Super 8x FOPFlash (TOPFlash mutant; Addgene plasmid # 12457), pCMV-Renilla ^32^, and one of the pcDNA3 plasmids encoding a peptide-guided uAb or control construct (e.g., CHIPΔTPR). Cells were transfected with a total of 100 ng of plasmid DNA in a ratio of TOPFlash/FOPFlash : Renilla : pcDNA3 = 1:0.1:3 using jetPRIME reagent. After 48 h of incubation, cells were lysed and the firefly and Renilla luminescence signals were measured sequentially by the dual-luciferase reporter kit (Promega). Luminescence was read on a microplate reader (Tecan Spark). All luciferase signals were measured and normalized against the control Renilla signals. For TOPFlash analysis of siRNA, an identical protocol was followed but with CTNNB1 siRNA at a final concentration of 100 nM per transfection instead of pcDNA3 plasmid.

### HiBiT assay

A total of 0.3 × 10^6^ DLD1 cells with CTNNB1-HiBiT CRISPR knock-in (Promega) were seeded per well in clear, flat 6-well plates. After 24 h incubation at 37 °C, 5% CO_2_, cells were transfected with 2 μg of pcDNA3 plasmid DNA encoding a uAb or control construct or with 100 nM of CTNNB1 siRNA using jetPRIME reagent. Cells were incubated for 48 h and then detached and harvested with 0.05% trypsin-EDTA. Cells were pelleted at 500 × g for 10 min and lysed, after which the cytosol, membrane, and nuclear fractions were isolated from lysates using a Subcellular Protein Fractionation Kit (ThermoFisher) per the manufacturer’s instructions. For HiBiT analysis, 20 μl of each fraction was added per well in white-bottom 96-well plates. 100 μl of the reagent from the Nano-Glo HiBiT Lytic Detection System kit (Promega) was added to each well followed by incubation for 30–60 min at room temperature. Luminescence was read on a plate reader (Tecan Spark). The total protein concentration in each well was measured by BCA assay. Signals for samples derived from uAb-expressing cells were normalized first by total protein concentration and then by the signals of samples derived from the cells only control.

### Real-time quantitative PCR analysis

DLD1 cells were seeded in a 6-well plate at a density of 2.5 × 10^5^ cells/well in 2 mL of RPMI1640 medium supplemented with 10% FBS, 24 h prior to transfection. On the day of transfection, DLD1 cells were transfected with 2 µg of pcDNA3 plasmid DNA encoding a uAb or control construct or with 100 nM of CTNNB1 siRNA using jetPRIME reagent, after which plates were incubated at 37 °C with 5% CO_2_. At 24 h post-transfection, cells were detached with 0.25% trypsin-EDTA and re-seeded in a 60 mm dish in 5 mL media. Cells were incubated at 37 °C, with 5% CO_2_, for 3 days. On the day of cell harvest, media was removed from the plate and 1 mL of TRIzol (ThermoFisher) was added to each well. RNA was extracted and purified according to the manufacturer’s instructions. RNA was converted to cDNA using the High-Capacity cDNA Reverse Transcription Kit (ThermoFisher). qPCR was performed with 25 ng of cDNA and SYBR Green Universal Master Mix (ThermoFisher) on a QuantStudio 7 Pro Real-Time PCR System. Cycle conditions were 95 °C for 10 min followed by 40 cycles of 95 °C for 20 sec and 60 °C for 1 min. Primers for each target gene were as follows: *AXIN2*: forward: TAACCCCTCAGAGCGATGGA, reverse: AGTTCCTCTCAGCAATCGGC; *CYP1A2*: forward: CCTTCGCTACCTGCCTAACC, reverse: CTCTAGGCCCCTTCTTGCTG; *c-Myc*: forward: CACCACCAGCAGCGACTCT; reverse: CTCTTGAGGACCAGTGGGCT; and *GAPDH*: forward: ATGGGGAAGGTGAAGGTCGG, reverse: TCCCGTTCTCAGCCTTGACG. All data were normalized to the CHIPΔTPR sample.

### Colony formation assay

DLD1 cells were seeded and transfected in the same manner as described above for the qPCR analysis. At 24 h post-transfection, cells were detached with 0.25% trypsin-EDTA and re-seeded in a 100 mm dish at a density of 2,000 cells/dish in 10 mL media. Cells were incubated at 37 °C, with 5% CO_2_, for 5 days. On the day of colony staining, media was removed from the dish, and cells as monolayers were gently rinsed two times with ample 1x PBS. A total of 5 mL of 0.5% (w/v) crystal violet (Sigma) was added to each dish, and the mixture was incubated for 15–20 min at room temperature. Crystal violet solution was removed, and culture dishes were rinsed four times with 8 mL of 1x PBS followed by imaging. Colony counting was performed by ImageJ using the same threshold for all samples. Data were normalized by colony counts of the CHIPΔTPR sample.

### MTS cell proliferation assay

A total of 5–7 × 10^4^ DLD1 cells were seeded the same way as for the TOPFlash assay but in clear, flat 96-well plates, one for each time point. After 24 h of incubation at 37 °C, with 5% CO_2_, cells were transfected with 100 ng of pcDNA3 plasmid DNA encoding a uAb or control construct or with 100 nM of CTNNB1 siRNA using jetPRIME reagent. Fresh media was replaced after 4–6 h of incubation. The plates were further incubated for 24 h and 48 h post transfection, and samples were collected at each of those time points. A total of 20 µL of MTS assay reagent (Promega) was added to each well, followed by another 1 h incubation. Absorbance was measured using a microplate reader (Tecan Spark) at a wavelength of 490 nm to assess cell viability and proliferation.

### Production of synthetic mRNA

Synthetic mRNAs corresponding to CHIPΔTPR, scr-CHIPΔTPR, and Ecad-30-CHIPΔTPR were produced using *in vitro* transcription (IVT). Briefly, mRNA sequences including a 5’ untranslated region and a 3’ untranslated region were each cloned in DNA plasmids downstream of an RNA promoter and upstream of a poly(A)120 region. The pDNA was expanded in *E. coli*, purified, and linearized downstream of the poly(A) region with BspQI. The linearized pDNA was combined with RNA polymerase, ATP, CTP, GTP, N1MePseudoUTP, and Mg^2+^ and incubated at 37 °C for 2 h. DNase I was added to the reaction to digest the pDNA template to stop the reaction and the RNA was purified. To cap the RNA, the purified RNA was combined with Vaccinia capping enzyme, *S*-adenosylmethionine, GTP, and Mg^2+^, incubated at 37 °C for 2 h, and purified. Purity of the mRNA was evaluated by capillary gel electrophoresis and content was evaluated by A260/280 absorbance spectroscopy.

### Preparation of mRNA-LNP formulations

Synthetic mRNAs were encapsulated in LNPs with Lipid 10 (Genevant Sciences) serving as the ionizable lipid in all formulations. This lipid was designed for i.v. delivery and was synthesized as described previously ^54^. All mRNAs were encapsulated in LNPs using a controlled mixing process (US 9005654) in which an aqueous solution of mRNA in acetate buffer at pH 5 was combined with an ethanolic solution of lipids in a T-shaped impingement zone. The lipid mix contained a PEG-conjugated lipid, Lipid 10 as the ionizable lipid, cholesterol, and DSPC at a molar ratio of 1.6:54.6:32.8:10.9, respectively, at a total lipid-to-RNA ratio of 20:1 weight/weight. Ethanol was removed by tangential flow ultrafiltration, followed by buffer exchange and concentration. The formulations were adjusted to 0.5 mg/mL and sterile filtered through a 0.2 μm PES membrane. Aliquots were subsequently stored frozen at –80 °C in Tris-sucrose buffer, pH 8.0, until the day of dosing.

### LNP characterization

LNP formulations were characterized by particle size analysis using a dynamic light scattering (DLS) instrument. Briefly, LNPs were diluted to 0.8–1.6 ng/μL total mRNA in PBS, pH 7.4 and transferred into a polystyrene cuvette to measure particle size and polydispersity by DLS (Malvern Nano ZS Zetasizer), using RI of 1.590 and absorption of 0.010 in PBS at 25 °C and viscosity of 0.9073 cP and refractive index (RI) of 1.332. Measurements were made with 10 s run durations with the number of runs automatically determined. Each measurement had a fixed position of 4.65 mm in the cuvette with an automatic attenuation selection. Diameters were reported as Z-average.

### *In vivo* administration of LNPs in mice

LNP formulations encapsulating mRNA corresponding to CHIPΔTPR, scr-CHIPΔTPR, and Ecad-30-CHIPΔTPR were administered intravenously by tail vein injection at a dose of 1.0 mg/kg of mRNA to female BALB/c mice (strain # 000651, 6–7 weeks old; Jackson Laboratory). On the day of injection, the LNP stocks were filtered and diluted to the required dosing concentration with PBS. At 1-day post-injection, animals were euthanized under carbon dioxide, and livers were harvested and collected. Livers were also collected from mice receiving CHIPΔTPR and Ecad-30-CHIPΔTPR at 3-, 5-, and 7-days post-injection. Harvested livers were weighed and flash frozen in liquid nitrogen and stored at –80 °C overnight. The next day, livers were homogenized to obtain lysates, which were subsequently fractionated into cytosolic and membrane fractions using a Subcellular Protein Fractionation Kit (ThermoFisher) per the manufacturer’s instructions. The isolated fractions were then subjected to immunoblotting analysis as described above. All animal experiments were reviewed and approved by the Institution of Animal Care and Use Committees (IACUC) of Cornell University under protocol # 2019-0063.

### Statistics and reproducibility

To ensure robust reproducibility of all results, experiments were performed with at least three biological replicates and at least three technical measurements. Sample sizes were not predetermined based on statistical methods but were chosen according to the standards of the field (at least three independent biological replicates for each condition), which gave sufficient statistics for the effect sizes of interest. All data were reported as average values with error bars representing standard deviation (SD). For individual samples, statistical significance was determined by paired Student’s *t* tests (\**p* < 0.05, \*\**p* < 0.01; \*\*\**p* < 0.001; \*\*\*\**p* < 0.0001) using Prism 10 for MacOS version 10.3.0. No data were excluded from the analyses. The experiments were not randomized. The investigators were not blinded to allocation during experiments and outcome assessment.

## Data Availability

All data generated or analyzed during this study are included in this article and its Supplementary Information/Source Data file that are provided with this paper.

## Code availability

SaLT&PepPr training data and SaLT&PepPr code can be found at: https://huggingface.co/ubiquitx/saltnpeppr, which includes an easy-to-use Colab notebook for peptide generation.

## Acknowledgements.

We thank UbiquiTx, Inc. for access to the SaLT&PepPr algorithm and the Duke Compute Cluster for providing the high-performance computing and database resources that have contributed to the research reported within this manuscript. We further thank Allison Chen for assistance with evaluating uAb-mRNA-LNP formulations and Dr. Sheng Zhang at the Cornell Proteomics and Metabolomics Core Facility for assistance with MS-based proteomics analysis. This work was supported by the National Institutes of Health (grants 1R41GM153081-01 to M.P.D and R21CA278468 to P.C.), the National Science Foundation (grant CBET-1605242 to M.P.D.), the Defense Threat Reduction Agency (grant HDTRA1-20-10004 to M.P.D.), and UbiquiTx, Inc. (through a sponsored research agreement to M.P.D.). A.A. and D.C. were supported by an NIH/NIGMS Chemical Biology Interface Training Grant (T32GM138826). C.D. was supported by a Hartwell Foundation Postdoctoral Fellowship. A.C. was supported by a National Science Foundation Graduate Research Fellowship.

## Author Contributions

T.Y. designed research, performed research, analyzed data, and wrote the paper. A.A., C.R., C.M., D.C., and C.D. designed research, performed research and analyzed data. L.H., S.V., and S.G. developed and implemented pLM models and analyzed data. K.L. and J.H. designed, performed and directed research. P.F., L.M.C., D.P., and C.A.A. designed and directed research. P.C. and M.P.D. designed and directed research, analyzed data, and wrote the paper. All authors read and approved the final manuscript.

## REFERENCES

1. Wodarz, A. & Nusse, R. Mechanisms of Wnt signaling in development. Annu Rev Cell Dev Biol 14, 59–88 (1998).

2. Peifer, M. & Polakis, P. Wnt signaling in oncogenesis and embryogenesis--a look outside the nucleus. Science 287, 1606–1609 (2000).

3. Nusse, R. & Clevers, H. Wnt/beta-Catenin Signaling, Disease, and Emerging Therapeutic Modalities. Cell 169, 985–999 (2017).

4. Buechel, D. et al. Parsing beta-catenin’s cell adhesion and Wnt signaling functions in malignant mammary tumor progression. Proc Natl Acad Sci U S A 118 (2021).

5. Bienz, M. β-catenin: a pivot between cell adhesion and Wnt signalling. Curr Biol 15, R64–67 (2005).

6. He, T.C. et al. Identification of c-MYC as a target of the APC pathway. Science 281, 1509–1512 (1998).

7. Tetsu, O. & McCormick, F. Beta-catenin regulates expression of cyclin D1 in colon carcinoma cells. Nature 398, 422–426 (1999).

8. Shtutman, M. et al. The cyclin D1 gene is a target of the beta-catenin/LEF-1 pathway. Proc Natl Acad Sci U S A 96, 5522–5527 (1999).

9. Moon, R.T., Kohn, A.D., De Ferrari, G.V. & Kaykas, A. WNT and beta-catenin signalling: diseases and therapies. Nat Rev Genet 5, 691–701 (2004).

10. Polakis, P. The many ways of Wnt in cancer. Curr Opin Genet Dev 17, 45–51 (2007).

11. Cui, C., Zhou, X., Zhang, W., Qu, Y. & Ke, X. Is β-catenin a druggable target for cancer therapy? Trends Biochem Sci 43, 623–634 (2018).

12. Dudek, H. et al. Knockdown of beta-catenin with dicer-substrate siRNAs reduces liver tumor burden in vivo. Mol Ther 22, 92–101 (2014).

13. Zeng, G., Apte, U., Cieply, B., Singh, S. & Monga, S.P. siRNA-mediated beta-catenin knockdown in human hepatoma cells results in decreased growth and survival. Neoplasia 9, 951–959 (2007).

14. Ganesh, S. et al. Direct pharmacological inhibition of β-catenin by RNA interference in tumors of diverse origin. Mol Cancer Ther 15, 2143–2154 (2016).

15. Conacci-Sorrell, M., Zhurinsky, J. & Ben-Ze’ev, A. The cadherin-catenin adhesion system in signaling and cancer. J Clin Invest 109, 987–991 (2002).

16. Gavrilov, K. et al. Enhancing potency of siRNA targeting fusion genes by optimization outside of target sequence. Proc Natl Acad Sci U S A 112, E6597–6605 (2015).

17. Kobayashi, Y., Tian, S. & Ui-Tei, K. The siRNA off-target effect Is determined by base-pairing stabilities of two different regions with opposite effects. Genes (Basel) 13 (2022).

18. Zhou, P. Targeted protein degradation. Curr Opin Chem Biol 9, 51–55 (2005).

19. Lopez-Barbosa, N., Ludwicki, M.B. & DeLisa, M.P. Proteome editing using engineered proteins that hijack cellular quality control machinery. AIChE J 66, e16854 (2020).

20. Tsai, J.M., Nowak, R.P., Ebert, B.L. & Fischer, E.S. Targeted protein degradation: from mechanisms to clinic. Nat Rev Mol Cell Biol 25, 740–757 (2024).

21. Simonetta, K.R. et al. Prospective discovery of small molecule enhancers of an E3 ligase-substrate interaction. Nat Commun 10, 1402 (2019).

22. Liao, H. et al. A PROTAC peptide induces durable beta-catenin degradation and suppresses Wnt-dependent intestinal cancer. Cell Discov 6, 35 (2020).

23. Portnoff, A.D., Stephens, E.A., Varner, J.D. & DeLisa, M.P. Ubiquibodies, synthetic E3 ubiquitin ligases endowed with unnatural substrate specificity for targeted protein silencing. J Biol Chem 289, 7844–7855 (2014).

24. Caussinus, E., Kanca, O. & Affolter, M. Fluorescent fusion protein knockout mediated by anti-GFP nanobody. Nat Struct Mol Biol 19, 117–121 (2011).

25. Lopez-Barbosa, N., Ludwicki, M.B. & DeLisa, M.P. Proteome editing using engineered proteins that hijack cellular quality control machinery. AIChE J 66, e16854 (2020).

26. Ludwicki, M.B. et al. Broad-spectrum proteome editing with an engineered bacterial ubiquitin ligase mimic. ACS Cent Sci 5, 852–866 (2019).

27. Pan, T. et al. A recombinant chimeric protein specifically induces mutant KRAS degradation and potently inhibits pancreatic tumor growth. Oncotarget 7, 44299–44309 (2016).

28. Poirson, J. et al. Proteome-scale discovery of protein degradation and stabilization effectors. Nature 628, 878–886 (2024).

29. Brixi, G., et al. SaLT&PepPr is an interface-predicting language model for designing peptide-guided protein degraders. Commun Biol 6, 1081 (2023).

30. Stephens, E.A. et al. Engineering single pan-specific ubiquibodies for targeted degradation of all forms of endogenous ERK protein kinase. ACS Synth Biol 10, 2396–2408 (2021).

31. Bery, N., Miller, A. & Rabbitts, T. A potent KRAS macromolecule degrader specifically targeting tumours with mutant KRAS. Nat Commun 11, 3233 (2020).

32. Cong, F., Zhang, J., Pao, W., Zhou, P. & Varmus, H. A protein knockdown strategy to study the function of beta-catenin in tumorigenesis. BMC Mol Biol 4, 10 (2003).

33. Su, Y., Ishikawa, S., Kojima, M. & Liu, B. Eradication of pathogenic beta-catenin by Skp1/Cullin/F box ubiquitination machinery. Proc Natl Acad Sci U S A 100, 12729–12734 (2003).

34. Teng, K.W. et al. Selective and noncovalent targeting of RAS mutants for inhibition and degradation. Nat Commun 12, 2656 (2021).

35. Roth, S. et al. Targeting endogenous K-RAS for degradation through the affinity-directed protein missile system. Cell Chem Biol 27, 1151–1163 e1156 (2020).

36. Lim, S. et al. Exquisitely specific anti-KRAS biodegraders inform on the cellular prevalence of nucleotide-loaded states. ACS Cent Sci 7, 274–291 (2021).

37. Bhat, S., et al. De novo design of peptide binders to conformationally diverse targets with contrastive language modeling. bioRxiv (2024).

38. Chatterjee, P. et al. Targeted intracellular degradation of SARS-CoV-2 via computationally optimized peptide fusions. Commun Biol 3, 715 (2020).

39. Chen, T. et al. PepMLM: Target sequence-conditioned generation of therapeutic peptide binders via span masked language modeling. ArXiv (2024).

40. Lin, Z. et al. Evolutionary-scale prediction of atomic-level protein structure with a language model. Science 379, 1123–1130 (2023).

41. Choi, H.J., Huber, A.H. & Weis, W.I. Thermodynamics of β-catenin-ligand interactions: the roles of the N- and C-terminal tails in modulating binding affinity. J Biol Chem 281, 1027–1038 (2006).

42. Huber, A.H. & Weis, W.I. The structure of the beta-catenin/E-cadherin complex and the molecular basis of diverse ligand recognition by beta-catenin. Cell 105, 391–402 (2001).

43. Wu, G. et al. Structure of a beta-TrCP1-Skp1-beta-catenin complex: destruction motif binding and lysine specificity of the SCF(beta-TrCP1) ubiquitin ligase. Mol Cell 11, 1445–1456 (2003).

44. Polakis, P. Wnt signaling in cancer. Cold Spring Harb Perspect Biol 4 (2012).

45. Veeman, M.T., Slusarski, D.C., Kaykas, A., Louie, S.H. & Moon, R.T. Zebrafish prickle, a modulator of noncanonical Wnt/Fz signaling, regulates gastrulation movements. Curr Biol 13, 680–685 (2003).

46. Traenkle, B. et al. Monitoring interactions and dynamics of endogenous beta-catenin with intracellular nanobodies in living cells. Mol Cell Proteomics 14, 707–723 (2015).

47. Qian, S.B. et al. Engineering a ubiquitin ligase reveals conformational flexibility required for ubiquitin transfer. J Biol Chem 284, 26797–26802 (2009).

48. Peng, J. et al. A proteomics approach to understanding protein ubiquitination. Nat Biotechnol 21, 921–926 (2003).

49. Jho, E.H. et al. Wnt/beta-catenin/Tcf signaling induces the transcription of Axin2, a negative regulator of the signaling pathway. Mol Cell Biol 22, 1172–1183 (2002).

50. Sekine, S., Lan, B.Y., Bedolli, M., Feng, S. & Hebrok, M. Liver-specific loss of beta-catenin blocks glutamine synthesis pathway activity and cytochrome p450 expression in mice. Hepatology 43, 817–825 (2006).

51. Baden, L.R. et al. Efficacy and Safety of the mRNA-1273 SARS-CoV-2 Vaccine. N Engl J Med 384, 403–416 (2021).

52. Polack, F.P. et al. Safety and Efficacy of the BNT162b2 mRNA Covid-19 Vaccine. N Engl J Med 383, 2603–2615 (2020).

53. Adams, D. et al. Patisiran, an RNAi Therapeutic, for Hereditary Transthyretin Amyloidosis. N Engl J Med 379, 11–21 (2018).

54. Lam, K. et al. Unsaturated, trialkyl ionizable lipids are versatile lipid-nanoparticle components for therapeutic and vaccine applications. Adv Mater 35, e2209624 (2023).

55. Kong, F. et al. Engineering a single ubiquitin ligase for the selective degradation of all activated ErbB receptor tyrosine kinases. Oncogene 33, 986–995 (2014).

56. Zhang, J., Zheng, N. & Zhou, P. Exploring the functional complexity of cellular proteins by protein knockout. Proc Natl Acad Sci U S A 100, 14127–14132 (2003).

57. VanDyke, D., Taylor, J.D., Kaeo, K.J., Hunt, J. & Spangler, J.B. Biologics-based degraders - an expanding toolkit for targeted-protein degradation. Curr Opin Biotechnol 78, 102807 (2022).

58. Stankiewicz, M., Nikolay, R., Rybin, V. & Mayer, M.P. CHIP participates in protein triage decisions by preferentially ubiquitinating Hsp70-bound substrates. FEBS J 277, 3353–3367 (2010).

59. Cyr, D.M., Hohfeld, J. & Patterson, C. Protein quality control: U-box-containing E3 ubiquitin ligases join the fold. Trends Biochem Sci 27, 368–375 (2002).

60. Lim, S. et al. bioPROTACs as versatile modulators of intracellular therapeutic targets including proliferating cell nuclear antigen (PCNA). Proc Natl Acad Sci U S A 117, 5791–5800 (2020).

61. Ma, Y. et al. Targeted degradation of KRAS by an engineered ubiquitin ligase suppresses pancreatic cancer cell growth in vitro and in vivo. Mol Cancer Ther 12, 286–294 (2013).

62. Hatakeyama, S., Watanabe, M., Fujii, Y. & Nakayama, K.I. Targeted destruction of c-Myc by an engineered ubiquitin ligase suppresses cell transformation and tumor formation. Cancer Res 65, 7874–7879 (2005).

63. Baltz, M.R., Stephens, E.A. & DeLisa, M.P. Design and functional characterization of synthetic E3 ubiquitin ligases for targeted protein depletion. Curr Protoc Chem Biol 10, 72–90 (2018).

64. Chen, T., Hong, L., Yudistyra, V., Vincoff, S. & Chatterjee, P. Generative design of therapeutics that bind and modulate protein states. Curr Opin Biomed Eng 28, 100496 (2023).

65. Mitragotri, S., Burke, P.A. & Langer, R. Overcoming the challenges in administering biopharmaceuticals: formulation and delivery strategies. Nat Rev Drug Discov 13, 655–672 (2014).

66. Chen, W., Lee, J., Cho, S.Y. & Fine, H.A. Proteasome-mediated destruction of the cyclin a/cyclin-dependent kinase 2 complex suppresses tumor cell growth in vitro and in vivo. Cancer Res 64, 3949–3957 (2004).

67. Cohen, J.C. et al. Transient in utero knockout (TIUKO) of C-MYC affects late lung and intestinal development in the mouse. BMC Dev Biol 4, 4 (2004).

68. Sufan, R.I. et al. Oxygen-independent degradation of HIF-alpha via bioengineered VHL tumour suppressor complex. EMBO Mol Med 1, 66–78 (2009).

69. Chan, A. et al. Lipid-mediated intracellular delivery of recombinant bioPROTACs for the rapid degradation of undruggable proteins. Nat Commun 15, 5808 (2024).

70. Pardi, N. et al. Expression kinetics of nucleoside-modified mRNA delivered in lipid nanoparticles to mice by various routes. J Control Release 217, 345–351 (2015).

71. Yamazaki, K. et al. Lipid nanoparticle-targeted mRNA formulation as a treatment for ornithine-transcarbamylase deficiency model mice. Mol Ther Nucleic Acids 33, 210–226 (2023).

72. Huang, X. et al. The landscape of mRNA nanomedicine. Nat Med 28, 2273–2287 (2022).

73. Schneider, C.A., Rasband, W.S. & Eliceiri, K.W. NIH Image to ImageJ: 25 years of image analysis. Nat Methods 9, 671–675 (2012).

