## Supplementary Information for "Programmable protein degraders enable selective knockdown of pathogenic β-catenin subpopulations *in vitro* and *in vivo*"

*To whom correspondence should be addressed:

### Supplementary Materials and Methods

**Proteasome inhibition with MG132.** To inhibit proteasomal degradation, DLD1 cells carrying empty plasmid or plasmid DNA encoding Ecad-30-CHIPΔTPR or CHIPΔTPR were transfected as described in the Materials and Methods section of the main paper. After 24 h of incubation, transfected cells were treated with either 5 μM MG132 or DMSO and incubated for an additional 24–48 h, after which cells were harvested for immunoblot analysis of cytosolic fractions.

**Protein expression and purification.** For expressing and purifying uAb degraders, each peptide-guided uAb or CHIPΔTPR construct was cloned into plasmid pET28a. The resulting plasmids were used to transform *E. coli* strain BL21(DE3). Plasmid-bearing cells were cultured overnight in Luria-Bertani (LB) media supplemented with appropriate antibiotics and the next day were diluted 1:100 in fresh LB media with appropriate antibiotics. When the optical density at 600 nm ($OD_{600}$) reached 0.4-0.6, cultures were induced with 1 mM isopropyl β-D-1-thiogalactopyranoside (IPTG) and further cultured at 30 °C overnight. The next day, cells were harvested by centrifugation at 4,500 × g for 25 min at 4 °C. The resulting pellets were resuspended in 10 mL of 1x phosphate-buffered saline (PBS) and lysed using an EmulsiFlex-C5 high-pressure homogenizer (Avestin). Lysates were cleared of insoluble material by centrifugation at 10,000 × g for 10 min at 4 °C. Clarified lysates containing polyhistidine (6x-His)-tagged proteins were subjected to gravity-flow $Ni^{2+}$-affinity purification using HisPur Ni-NTA resin (ThermoFisher) following the manufacturer's protocols. Purified proteins were desalted in 1x PBS, and the final purity of all proteins was confirmed by Coomassie-blue staining of SDS-PAGE gels.

**ELISA.** 96-well plates (MaxiSorp; Nunc Nalgene) were incubated with 4 μg/mL of purified β-catenin (Biomatik) diluted in 1x PBS, pH 7.4, 80 μL/well, at 4 °C overnight. Plates were incubated with 200 μL blocking buffer containing 3% (w/v) BSA in 1x TBST overnight at 4°C. Plates were washed three times by adding 200 μL of 1x PBST (PBS, 0.1% (v/v) Tween 20). All uAbs were biotinylated using the EZ-Link Sulfo-NHS-LC-Biotin (Thermo) biotinylation reagent, followed by desalting with Zeba spin columns (Thermo) according to the manufacturer's protocol. Biotinylation was verified by dot blots probed with Streptavidin-HRP (Abcam, cat # ab7403; diluted 1:5000). Next, biotinylated uAbs were serially diluted in triplicate in 1x PBS and added to the ELISA plates for 1 h at 37 °C.

Plates were washed three times with 1x PBS-T. Plates were then incubated with HRP-conjugated streptavidin (ThermoFisher, cat # N100; diluted 1:20,000) for 1 h at room temperature with gentle shaking. After another three washes with 1x PBS-T, 100 µL of 3,3'-5,5'-tetramethylbenzidine substrate (1-Step Ultra TMB-ELISA; ThermoFisher) was added to each well, and the plates were incubated at room temperature in the dark. The reactions were stopped by addition of 100 µL of 2 M $H_2SO_4$ to each well, after which the absorbance of the plates was measured at a wavelength of 450 nm using a microplate reader (Tecan Spark).

**Biolayer interferometry analysis.** The dissociation constant for the purified uAbs to β-catenin was measured by biolayer interferometry (BLI) using an Octet RH16 instrument (Sartorious) in combination with streptavidin (SA) biosensor tips (Sartorius). Briefly, kinetic analysis was performed as follows: (1) blocking of biosensor tips by immersion for 30 s in Octet buffer (1x PBS containing 0.2 µm filtered 0.1% BSA and 0.05% (v/v) Tween 20); (2) uAb immobilization on biosensors by immersion of tips for 60 s in Octet buffer supplemented with 1 µg/ml biotinylated-uAbs prepared as described above; (3) wash of biosensor tips by immersion for 60 s in Octet buffer; (4) β-catenin association by immersion of treated tips for 300 s in Octet buffer containing purified β-catenin (Biomatik) at the following concentrations: 1480 nM, 740 nM, 370 nM, 185 nM, 92.5 nM, and 46.3 nM; and (5) β-catenin dissociation by immersion for 300 s in Octet buffer. All steps were performed at 30 °C. The kinetic data was analyzed using Octet Analysis Studio software v12.2.2.26 (Sartorius).

***In vitro* ubiquitination assays and β-catenin immunoprecipitation.** *In vitro* ubiquitination of recombinant β-catenin was done as previously described [1]. Briefly, 0.1 µM purified human UBE1 (R&D Systems), 4 µM human UbcH5α/UBE2D1 (R&D Systems), 3 µM purified uAb or CHIPΔTPR, 1.5 µM β-catenin (Origene) and 50 µM human ubiquitin (R&D Systems), were used in buffer containing 4 mM ATP, and 1 mM DTT in 20 mM MOPs, 100 mM KCl, 5 mM $MgCl_2$, pH 7.2. Reactions were carried out at 37 °C for 2 h (or collected in intervals of 30 min). β-catenin was immunoprecipitated from the reaction mixture using BC13-coated Dynabeads, which were generated in-house. Aliquots of the reaction mixture were taken at each time point and quenched by adding a solution of 200 mM EDTA pH 7 to a final concentration of 20 mM. Each sample was diluted in 100 mM sodium phosphate buffer pH 7 and mixed with 50 µL of BC13-coated Dynabeads (1 mg) with end-over-end rotation at 4 °C overnight. The next day, beads were pulled down over a magnet and washed three times with 1x PBS containing 0.05% (v/v) Tween-20 before being eluted by boiling in the presence of 50 µL 4x SDS loading dye. Samples were analyzed by immunoblotting using anti-β-catenin (Cell Signaling, cat # 8480S; diluted 1:1000) and anti-ubiquitin (Cell Signaling, cat # P4D1 3936; diluted 1:1,000) antibodies.

**Mass spectrometry analysis.** For LC-ESI-MS/MS analysis of ubiquitination, the *in vitro* ubiquitination reaction of β-catenin was performed as described above and the entire reaction mixture (~1 µg protein) was quenched by boiling in SDS loading dye. The mixture was resolved by SDS-PAGE, and protein bands were visualized by staining with Coomassie R-250 stain. The band of interest was then excised, and the excised gel band was cut into ~1 mm cubes and subjected to in-gel trypsin digestion. The excised gel pieces were washed/incubated at room temperature consecutively with 150 µL deionized water for 5 min, followed by 150 µl 50 mM ammonium bicarbonate in water/50% acetonitrile (ACN) for 10 min and finally 75 µl 100% ACN for 5 min. The dehydrated gel pieces were dried in a speed vacuum (SpeedVac SC110 Thermo Savant) and reduced with 40 µL of 10 mM dithiothreitol (DTT) in 100 mM ammonium bicarbonate in water for 1 h at 60 °C, then alkylated by adding 40 µl of 55 mM iodoacetamide in 100 mM ammonium bicarbonate (w/v) and incubation at room temperature, in the dark, for 45 min. Wash steps were repeated as described above. The gel pieces were dried in a speed vacuum and rehydrated with 40 µl trypsin (Promega Sequencing Grade) at 10 ng/µl in 50 mM ammonium bicarbonate/10% ACN on ice for 20 min, topped with 20 µl 50 mM ammonium bicarbonate in water, and incubated at 37 °C for 16 h. The digestion was stopped by addition of 60 µl 2% formic acid (FA) in water, incubated at room temperature for 10 min and the supernatant transferred to a clean polypropylene low-bind microfuge tube. The gel pieces were further extracted twice by adding 125 µl of 50% ACN/5% FA and vortexing at 1500 rpm for 10 min followed by sonication for 5 min, and once by adding 50 µl of 90% ACN/5% FA with incubation at room temperature for 5 min. All supernatants were combined in the corresponding microfuge tube, dried in a speed vacuum and redried from 50 µl water. The final sample was reconstituted in 2% ACN/0.5% FA and filtered through a 0.22 µm cellulose acetate spin filter (Corning Costar Spin-X) prior to nanoLC-MS/MS analysis.

**Protein Identification by nano LC/MS/MS Analysis.** The digests were reconstituted in 2% ACN with 0.5% FA for nanoLC-ESI-MS/MS analysis. The analysis was carried out using an Orbitrap Fusion™ Tribrid™ (ThermoFisher) mass spectrometer equipped with a nanospray Flex Ion Source and coupled with a Dionex UltiMate 3000 RSLCnano system (ThermoFisher) [2, 3]. The peptide samples (5 µL) were injected onto a PepMap C-18 RP viper trapping column (5 µm, 100 µm i.d x 20 mm) at 20 µL/min flow rate for rapid sample loading and then separated on a PepMap C-18 RP nano column (2 µm, 75 µm x 25 cm) at 35 °C. The tryptic peptides were eluted in a 90-min gradient of 5% to 33% ACN in 0.1% formic acid at 300 nL/min, followed by an 8-min ramping to 90% ACN-0.1% FA and an 8-min hold at 90% ACN-0.1% FA. The column was re-equilibrated with 0.1% FA for 25 min prior to the next run. The Orbitrap Fusion was operated in positive ion mode with spray voltage set at 1.5 kV and source temperature at 275 °C. External calibration for FT, IT and quadrupole mass analyzers was performed. In data-dependent acquisition (DDA) analysis, the instrument was operated using FT mass analyzer in MS scan to select precursor ions followed by 3-s "Top Speed" data-dependent CID ion trap MS/MS scans at 1.6 m/z quadrupole isolation for precursor peptides with multiple charged ions above a threshold ion count of 10,000 and normalized collision energy of 30%. MS survey scans at a resolving power of 120,000 (fwhm at *m*/*z* 200), for the mass range of *m*/*z* 375-1600. Dynamic exclusion parameters were set at 35 s of exclusion duration with ±10 ppm exclusion mass width. All data were acquired under Xcalibur 4.3 operation software (ThermoFisher).

**Data analysis.** The DDA raw files with MS and MS/MS were subjected to database searches using Proteome Discoverer (PD) 2.4 software (ThermoFisher) with the Sequest HT algorithm. The PD 2.4 processing workflow containing an additional node of Minora Feature Detector for precursor ion-based quantification was used for protein identification and relative quantitation of identified peptides and their modified forms. The database search was conducted against Homo Sapiens Uniprot database, which contains 20,359 sequences. The peptide precursor tolerance was set to 10 ppm, and fragment ion tolerance was set to 0.6 Da. Oxidation of M, deamidation of N and Q, phosphorylation on

S/T/Y, acetylation on K, ubiquitination on K, glycation on K/R were specified as dynamic modifications of amino acid residues; protein N-terminal acetylation, M-loss and M-loss plus acetylation were set as a variable modification; carbamidomethyl C was specified as a static modification. Only high-confidence peptides defined by Sequest HT with a 1% FDR by Percolator were considered for confident peptide identification.

**Occupancy rate calculation for ubiquitination.** The extraction ion chromatograms (XICs) for both ubiquitinated peptides and their counterpart native ones were obtained, and the peak areas were integrated using Xcalibur software. The modification occupancy rate was calculated by dividing the peak area of the modified peptide by the sum of the peak areas for both the native and modified peptide forms under assumption of similar ionization efficiencies for the different forms of the same peptide. All charge states of the modified and native peptides were combined and considered as a single form.

**LNP-mRNA delivery in Hep3B cells.** The human HCC cell line, Hep3B, was purchased from ATCC (cat # HB-8064). Hep3B cells were cultured in MEM media (ThermoFisher) supplemented with 100 units/mL penicillin, 100 mg/mL streptomycin, and 10% FBS (Corning) at 37 °C with 5% $CO_2$. Plasmid transfections, immunoblotting analysis, and TOPFlash assays were performed essentially as described in the Materials and Methods section of the main paper. For immunoblotting, $3 \times 10^5$ cells were seeded in a 6-well plate and 6 µg of mRNA-LNP formulations were used for transfection. For TOPFlash assays, $1 \times 10^4$ Hep3B cells were seeded in a 96-well plate. After 24 h, 10, 50, 75, 100, 150, or 200 ng of mRNA-LNP formulations were added to each well. Plates were incubated at 37 °C, with 5% $CO_2$ for 48 h before collecting samples for luciferase analysis.

### References


1. Stephens, E.A. et al. Engineering single pan-specific ubiquibodies for targeted degradation of all forms of endogenous ERK protein kinase. *ACS Synth Biol* **10**, 2396-2408 (2021).
2. Yang, Y., Thannhauser, T.W., Li, L. & Zhang, S. Development of an integrated approach for evaluation of 2-D gel image analysis: impact of multiple proteins in single spots on comparative proteomics in conventional 2-D gel/MALDI workflow. *Electrophoresis* **28**, 2080-2094 (2007).

3. Yang, Y., Anderson, E. & Zhang, S. Evaluation of six sample preparation procedures for qualitative and quantitative proteomics analysis of milk fat globule membrane. *Electrophoresis* **39**, 2332-2339 (2018).

**Supplementary Table 1.** SaLT&PepPr (SnP)-derived peptide sequences and scores based on interaction between β-catenin and the cytoplasmic domain of E-cadherin

| Peptide name | Peptide sequence | SnP probability score* |
|---|---|---|
| Ecad-1 | YDSLLVFDYE | 0.5114 |
| Ecad-2 | PPYDSLLVFDY | 0.5086 |
| Ecad-3 | TAPPYDSLLVFD | 0.5120 |
| Ecad-4 | TAPPYDSLLVFDYE | 0.5216 |
| Ecad-5 | APPYDSLLVFDYEGS | 0.5149 |
| Ecad-6 | PTAPPYDSLLVFDYEGSG | 0.5371 |
| Ecad-7 | DPTAPPYDSLLVFDYE | 0.5086 |
| Ecad-8 | PTAPPYDSLLVFDYEGS | 0.5349 |
| Ecad-9 | DPTAPPYDSLLVFDYEGS | 0.5312 |
| Ecad-10 | ADSDPTAPPYDSLLVFDYE | 0.5047 |
| Ecad-11 | AADSDPTAPPYDSLLVFDYE | 0.5052 |
| Ecad-12 | ADSDPTAPPYDSLLVFDYEGS | 0.5006 |
| Ecad-13 | LKAADSDPTAPPYDSLLVFDYE | 0.5130 |
| Ecad-14 | NLKAADSDPTAPPYDSLLVFDYE | 0.5175 |
| Ecad-15 | SDPTAPPYDSLLV | 0.4932 |
| Ecad-16 | PYDSLLVFDYE | 0.5217 |
| Ecad-17 | DSLLVFDYEG | 0.5131 |
| Ecad-18 | YDSLLVFDYEG | 0.5145 |
| Ecad-19 | PPYDSLLVFDYE | 0.5114 |
| Ecad-20 | PYDSLLVFDYEGS | 0.5290 |
| Ecad-21 | TAPPYDSLLVFDY | 0.5166 |
| Ecad-22 | APPYDSLLVFDYE | 0.5193 |
| Ecad-23 | PPYDSLLVFDYEG | 0.5177 |
| Ecad-24 | APPYDSLLVFDYEG | 0.5264 |
| Ecad-25 | ADTDPTAPPYDSLLV | 0.5297 |
| Ecad-26 | PTAPPYDSLLVFDYE | 0.5127 |
| Ecad-27 | DTDPTAPPYDSLLVFDYE | 0.5368 |
| Ecad-28 | ADTDPTAPPYDSLLVFDY | 0.5352 |
| Ecad-29 | DPTAPPYDSLLVFDYEGSGS | 0.5308 |
| Ecad-30 | PTAPPYDSLLVFDYEG | 0.5123 |
| Ecad-31 | TDPTAPPYDSLLVFDYEGS | 0.5284 |
| Ecad-32 | DPTAPPYDSLLVFDYEG | 0.5320 |
| Ecad-33 | DYEGSGSEAASLSSLNSSESDKDQ | 0.5088 |

***Note:** Peptides were down-selected by inputting specific domains of the target proteins into the SnP algorithm, which differs from our previous work which used the entire target protein sequence (Brixi et al. *Commun Biol* 2023). For E-cadherin, specifically, we input the cytoplasmic domain into the SnP algorithm, leading to cumulative binding site probability scores for each peptide.

**Supplementary Table 2.** SnP-derived peptide sequences and scores based on interaction between β-catenin and the WD40-repeat domains of β-TrCP

| Peptide name | Peptide sequence | SnP probability score* |
|---|---|---|
| β-TrCP-1 | GHRAAVNVVD | 0.3142 |
| β-TrCP-2 | RCIRFDNKRIVSGAY | 0.2381 |
| β-TrCP-3 | LVRCIRFDNKRIVSGAY | 0.2460 |
| β-TrCP-4 | AAVNVVDFDDKYIVSASG | 0.1936 |
| β-TrCP-5 | RAAVNVVDFDDKYIVSASG | 0.2174 |
| β-TrCP-6 | HRAAVNVVDFDDKYIVSASG | 0.2346 |
| β-TrCP-7 | HRAAVNVVDF | 0.3095 |
| β-TrCP-8 | AVNVVDFDDKYIVSASG | 0.2050 |
| β-TrCP-9 | HEELVRCIRF | 0.3047 |
| β-TrCP-10 | EELVRCIRFD | 0.3032 |
| β-TrCP-11 | VRCIRFDNKRIVSGAY | 0.2295 |

***Note:** Peptides were down-selected by inputting the WD40-repeat domains of β-TrCP into the SnP algorithm, leading to cumulative binding site probability scores for each peptide.

**Supplementary Table 3.** Interacting partner sequences input to SaLT&PepPr for identification of peptide sequences that interact with β-catenin.

| **Interacting partner name** | **Sequence** | **Uniport ID** |
|---|---|---|
| Cytosolic domain of E-cadherin (Ecad-CD) | RRRTVVKEPLLPPDDDTRDNVYYYDEEGGGEEDQDFDL SQLHRGLDARPEVTRNDVAPTLMSVPQYRPRPANPDEI GNFIDENLKAADSDPTAPPYDSLLVFDYEGSGSEAASLS SLNSSESDQDQDYDYLNEWGNRFKKLADMYGGGEDD | CADH1_Mouse |
| | LRRRAVVKEPLLPPEDDTRDNVYYYDEEGGGEEDQDFD LSQLHRGLDARPEVTRNDVAPTLMSVPRYLPRPANPDEI GNFIDENLKAADTDPTAPPYDSLLVFDYEGSGSEAASLS SLNSSESDKDQDYDYLNEWGNRFKKLADMYGGGEDD | CADH1_Human |
| β-TrCP (amino acids 301-590) | ETSKGVYCLQYDDQKIVSGLRDNTIKIWDKSTLECKRILT GHTGSVLCLQYDERVIITGSSDSTVRVWDVNAGEMLNTL IHHCEAVLHLRFNNGMMVTCSKDRSIAVWDMASPTDITL RRVLVGHRAAVNVVDFDDKYIVSASGDRTIKVWNTSTCE FVRTLNGHKRGIACLQYRDRLVVSGSSDNTIRLWDIECG ACLRVLEGHEELVRCIRFDNKRIVSGAYDGKIKVWDLMA ALDPRAPAGTLCLRTLVEHSGRVFRLQFDEFQIVSSSHD DTILIWDFLNDPAAHA | FBW1A_Mouse |
| | ETSKGVYCLQYDDQKIVSGLRDNTIKIWDKNTLECKRILT GHTGSVLCLQYDERVIITGSSDSTVRVWDVNTGEMLNTL IHHCEAVLHLRFNNGMMVTCSKDRSIAVWDMASPTDITL RRVLVGHRAAVNVVDFDDKYIVSASGDRTIKVWNTSTCE FVRTLNGHKRGIACLQYRDRLVVSGSSDNTIRLWDIECG ACLRVLEGHEELVRCIRFDNKRIVSGAYDGKIKVWDLVAA LDPRAPAGTLCLRTLVEHSGRVFRLQFDEFQIVSSSHDD TILIWDFLNDPAAQA | FBW1A_Human |

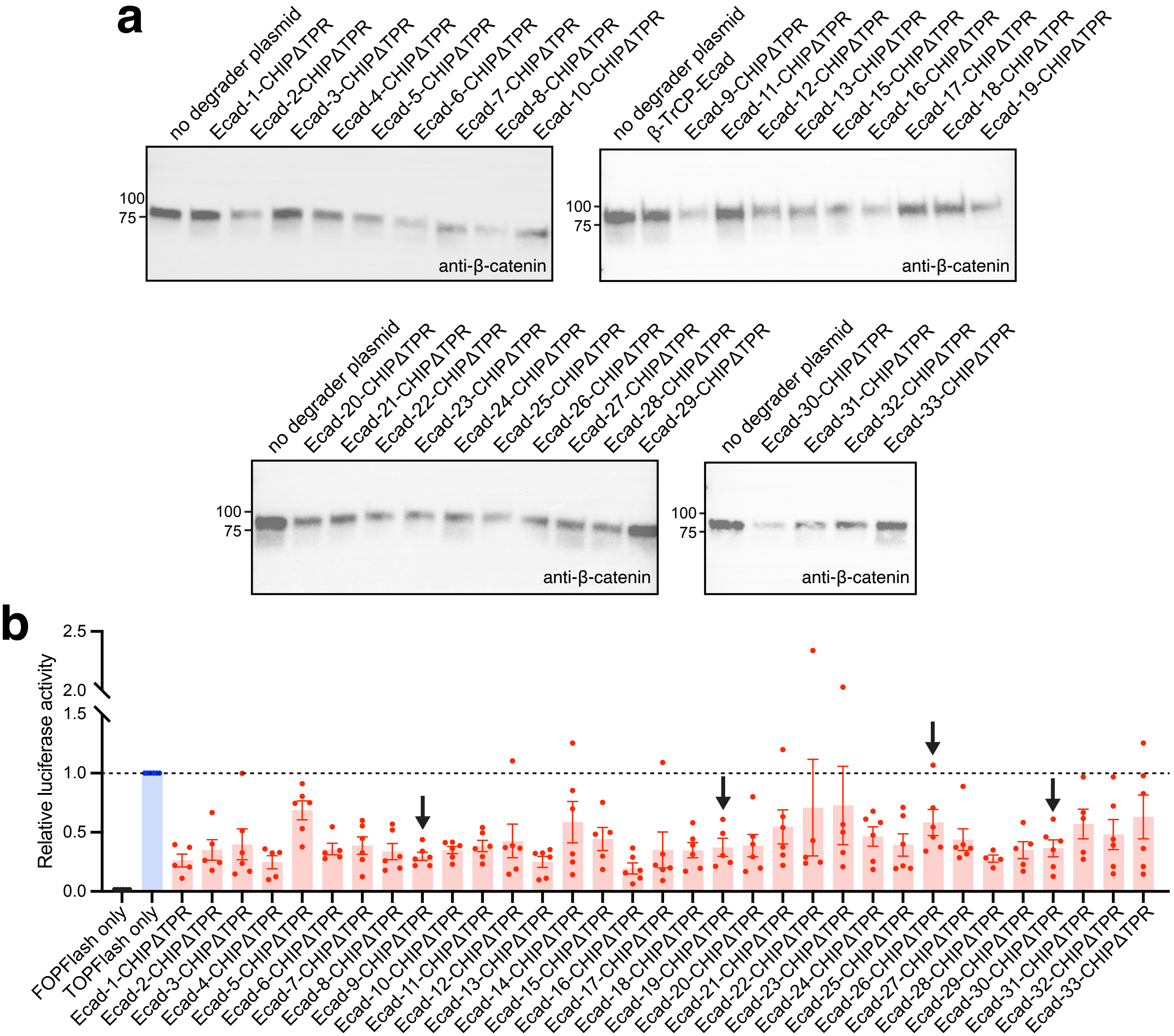


**Supplementary Figure 1. Cytosolic β-catenin knockdown by uAbs composed of E-cadherin-based guides.** (a) Immunoblot analysis of cytosolic β-catenin levels in DLD1 cells transfected with plasmid pcDNA3 encoding fusions between different E-cadherin-derived guide peptides and CHIPΔTPR. Also included is β-TrCP-Ecad degrader construct (top right blot) from Cong et al. (*BMC Mol Biol* 2003). Cells were harvested 48 h post transfection, after which cytoplasmic fractions were prepared from cell extracts and subjected to immunoblotting with anti-β-catenin antibody. Lanes were normalized by total protein content and molecular weight ($M_W$) markers are indicated at left. Blots are representative of at least three biological replicates. (b) β-catenin signaling activity in DLD1 cells co-transfected with TOPFlash reporter plasmid along with degrader-encoding plasmid from (a). Cells receiving only the FOPFlash or TOPFlash plasmid served as negative and positive controls, respectively. Luciferase signals in each sample were normalized to those measured in control cells receiving no degrader plasmid. Data are mean of at least three biological replicates ($n$ = 4−6) ± SD. Black arrows indicate uAbs that were downselected for further characterization.

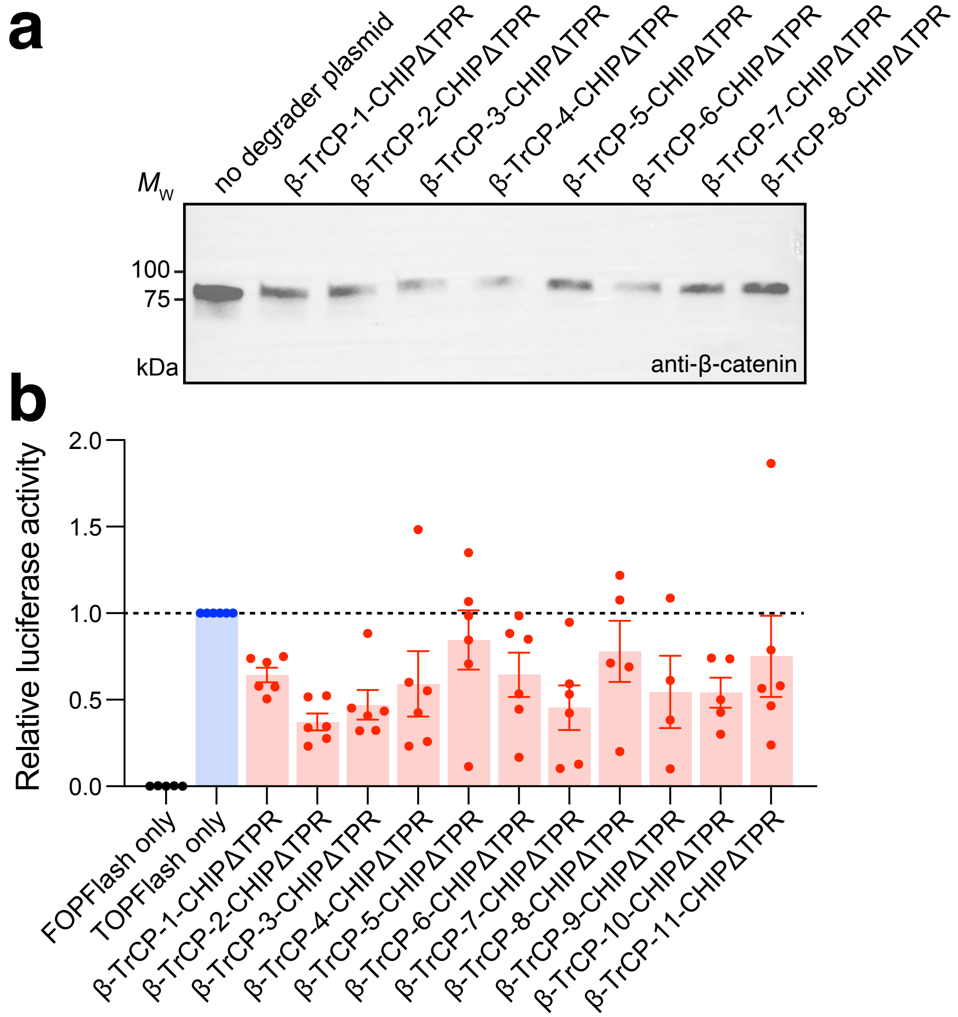


**Supplementary Figure 2. Cytosolic β-catenin knockdown by uAbs composed of β-TrCP-based guide peptides.** (a) Immunoblot analysis of cytosolic β-catenin levels in DLD1 cells transfected with plasmid pcDNA3 encoding fusions between different β-TrCP-derived guide peptides and CHIPΔTPR. Cells were harvested 48 h post-transfection, after which cytoplasmic fractions were prepared from cell extracts and subjected to immunoblotting with anti-β-catenin antibody. Lanes were normalized by total protein content and molecular weight ($M_w$) markers are indicated at left. Blots are representative of at least three biological replicates. (c) β-catenin signaling activity in DLD1 cells co-transfected with superTOPFlash reporter plasmid along with degrader-encoding plasmid from (b). Cells receiving only the FOPFlash or TOPFlash plasmid served as negative and positive controls, respectively. Luciferase signals in each sample were normalized to those measured in control cells receiving no degrader plasmid. Data are mean of at least three biological replicates ($n$ = 4−6) ± SD.

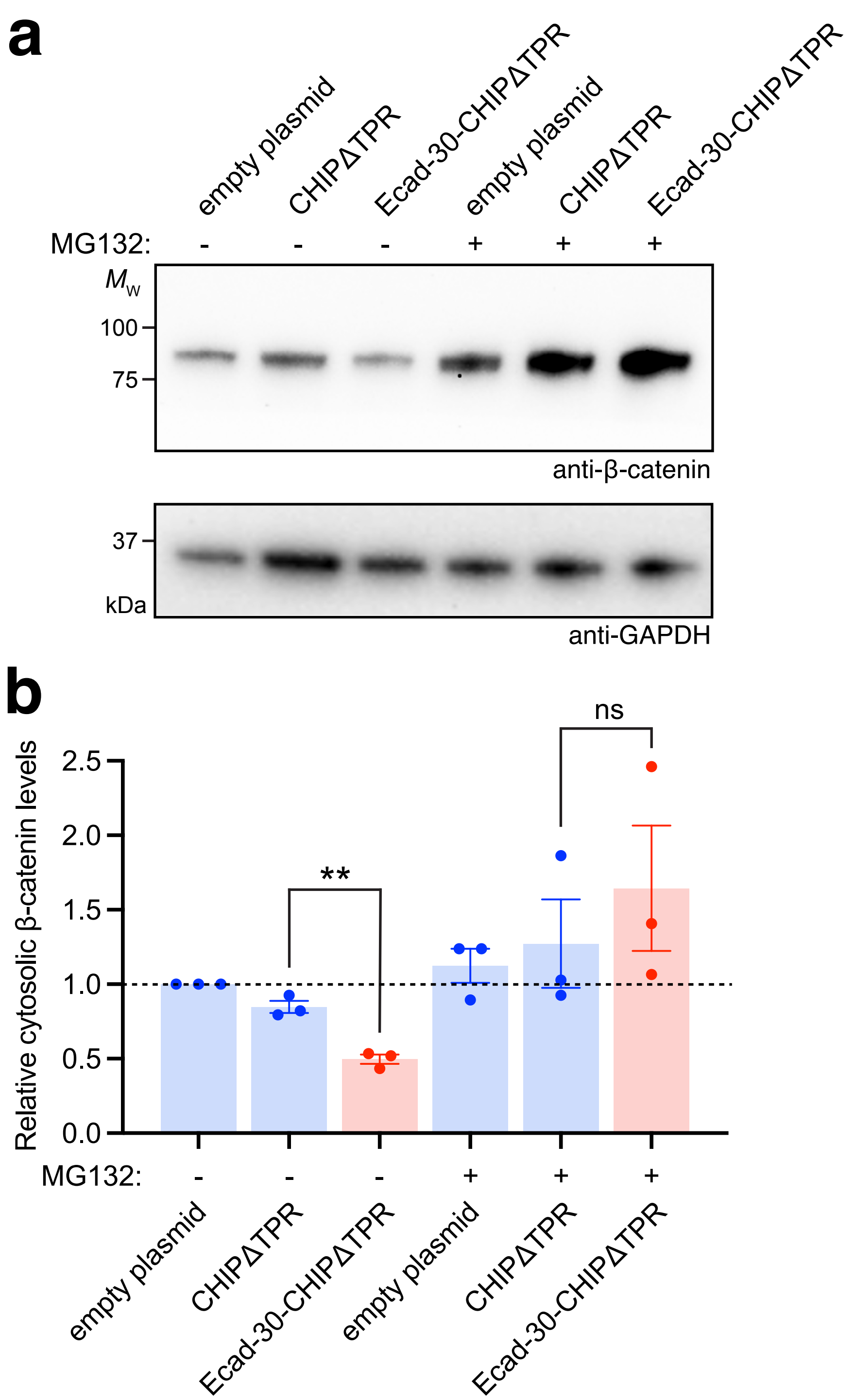


**Supplementary Figure 3. Proteasome-dependent β-catenin knockdown by Ecad-30-CHIPΔTPR degrader.** (a) Immunoblot analysis of cytosolic β-catenin levels in DLD1 cells transfected with empty plasmid pcDNA3 or pcDNA3 encoding either CHIPΔTPR or Ecad-30-CHIPΔTPR. MG132-treated (+) and untreated (-) cells were harvested 48 h post-transfection, after which cytoplasmic fractions were prepared from cell extracts and subjected to immunoblotting with anti-β-catenin antibody (top) and anti-GAPDH antibody (bottom), the latter serving as a loading control. Lanes were normalized by total protein content and molecular weight ($M_W$) markers are indicated at left. Blots are representative of three biological replicates. (b) Quantification of cytosolic β-catenin levels by densitometry analysis of immunoblots in panel (a). Band intensity was determined using ImageJ software with all β-catenin band intensities normalized to corresponding GAPDH band intensities. Relative β-catenin levels were then calculated by normalizing Ecad-30-CHIPΔTPR values to CHIPΔTPR values. Data are mean of biological replicates ($n$ = 3) ± SD. Statistical significance was determined by unpaired two-tailed Student's $t$-test. Calculated $p$ values are represented as follows: **, $p < 0.01$; ns, not significant.

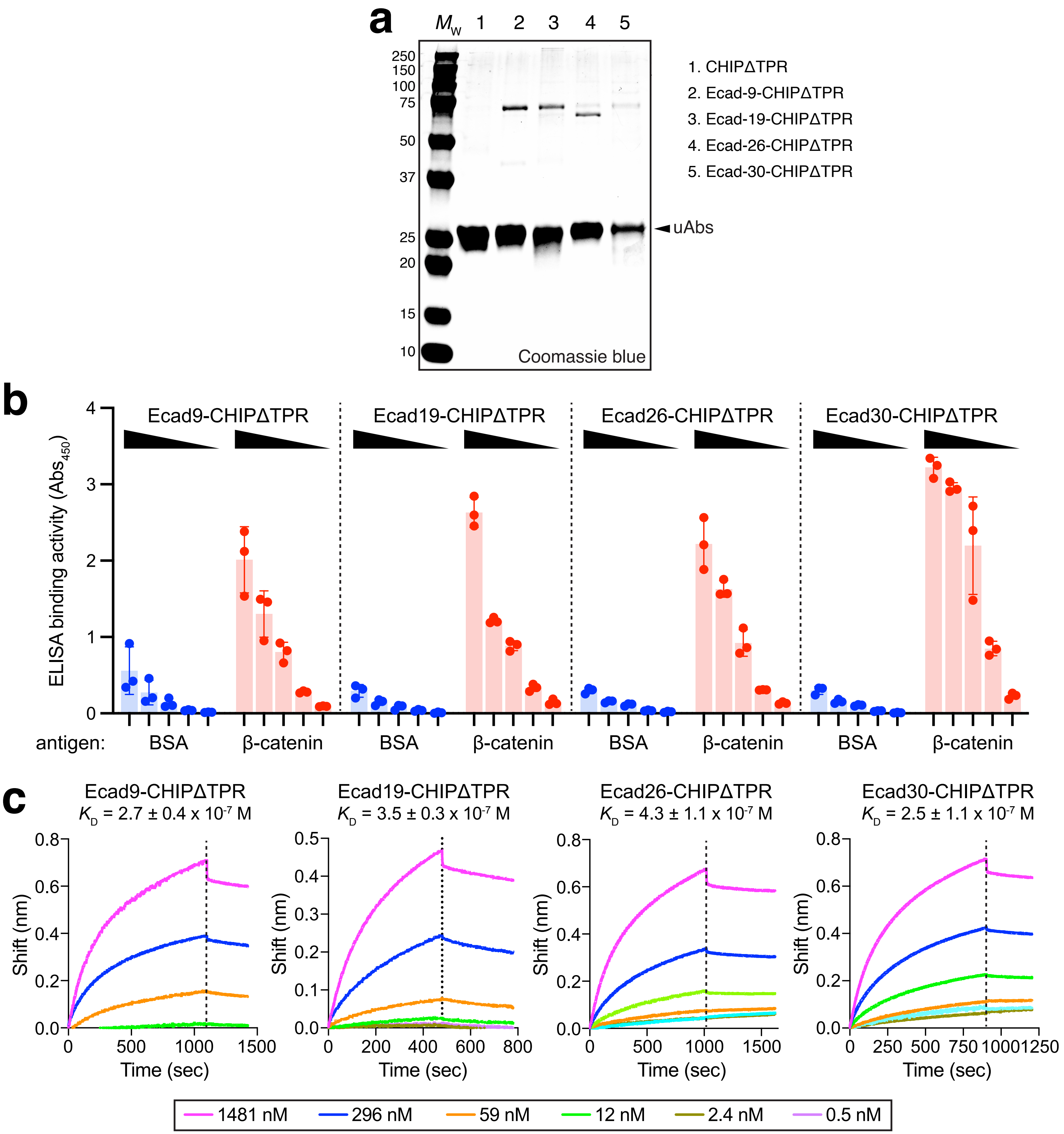


**Supplementary Figure 4. *In vitro* binding of β-catenin by peptide-guided uAbs.** (a) Coomassie blue-stained SDS-PAGE gel analysis of Ni-NTA-purified CHIPΔTPR and peptide-guided derivatives as indicated. A total of 2 µg of purified protein were loaded in each lane. Molecular weight ($M_W$) marker is shown at left. (b) ELISA analysis of purified uAbs with β-catenin or bovine serum albumin (BSA) as immobilized antigen. Increasing amounts of each uAb (851, 426, 213, 53, and 13 nM) were added to each well as indicated by black arrows. Data are mean of biological replicates ($n$ = 3) ± SD. (b) Biolayer interferometry (BLI) analysis of purified uAbs in (a) to quantify equilibrium binding constants ($K_D$) for the interaction between each uAb and β-catenin. Biotinylated uAbs were immobilized on streptavidin (SA)-coated sensors and subsequently used to bind β-catenin in solution. Response data are representative of replicate BLI experiments ($n$ = 2).

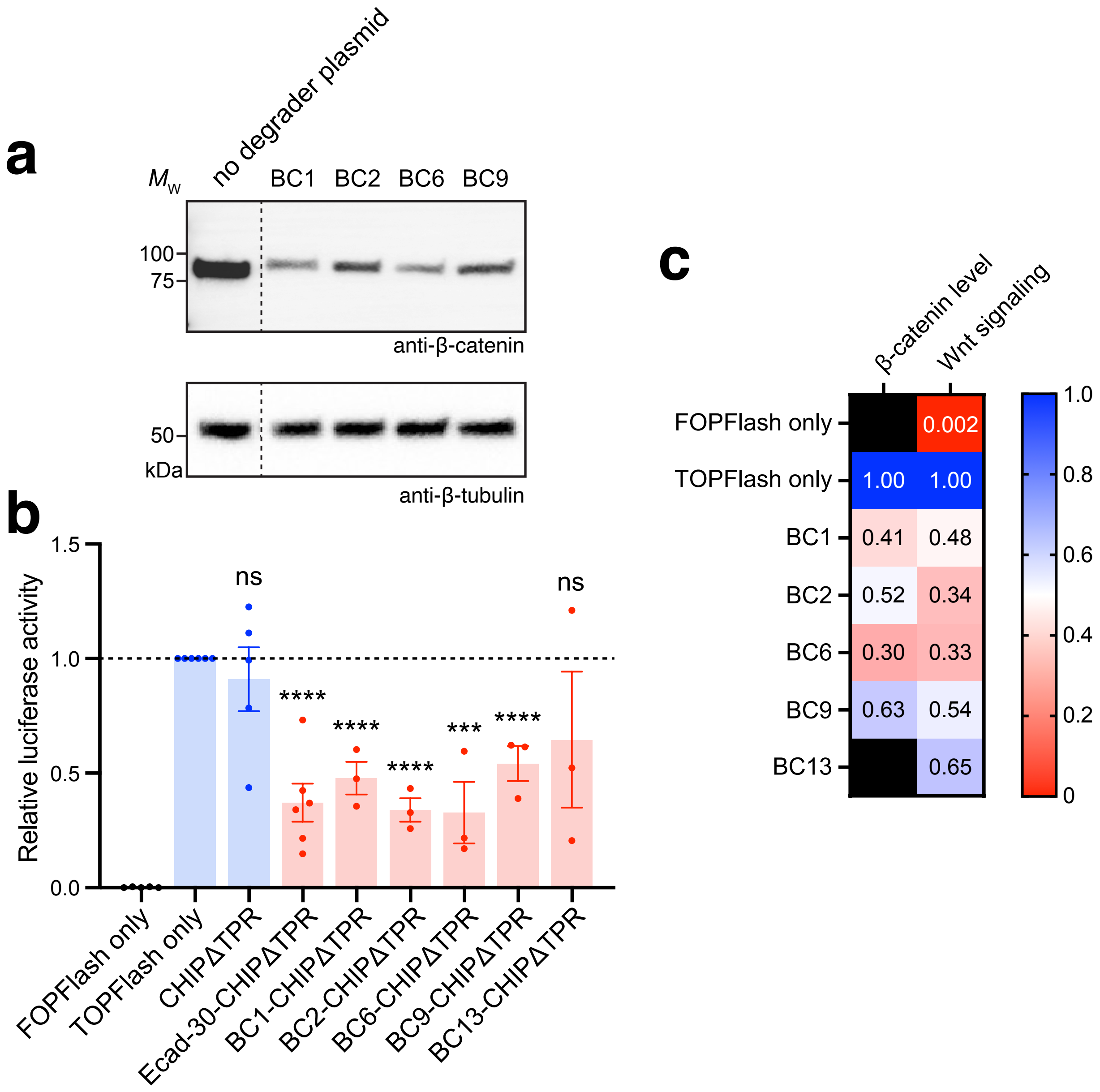


**Supplementary Figure 5. Cytosolic β-catenin knockdown by VHH-guided degraders.** (a) Immunoblot analysis of cytosolic β-catenin levels in DLD1 cells transfected with pcDNA3 encoding fusions between different β-catenin-specific VHHs and CHIPΔTPR. Cells were harvested 48 h post-transfection, after which cytoplasmic fractions were prepared from cell extracts and subjected to immunoblotting with anti-β-catenin antibody (top) and anti-β-tubulin antibody (bottom), the latter serving as a loading control. Lanes were normalized by total protein content and molecular weight ($M_w$) markers are indicated at left. Blot is representative of three biological replicates. Dashed line indicates splicing of same immunoblot to remove unused lanes. (b) β-catenin signaling activity in DLD1 cells co-transfected with TOPFlash reporter plasmid along with degrader-encoding plasmid from (a) or plasmid encoding Ecad-30-CHIPΔTPR construct. Cells receiving only the FOPFlash or TOPFlash plasmid served as negative and positive controls, respectively. Luciferase signals in each sample were normalized to those measured in control cells receiving no degrader plasmid. Data are mean of at least three biological replicates ($n$ = 3−6) ± SD. Statistical significance was determined by unpaired two-tailed Student's $t$-test. Calculated $p$ values are represented as follows: ***, $p < 0.001$; ****, $p < 0.0001$; ns, not significant. (c) Heatmap of normalized β-catenin levels determined by densitometry analysis of blot in (a) and normalized β-catenin signaling reported in (b). Black box = not tested.

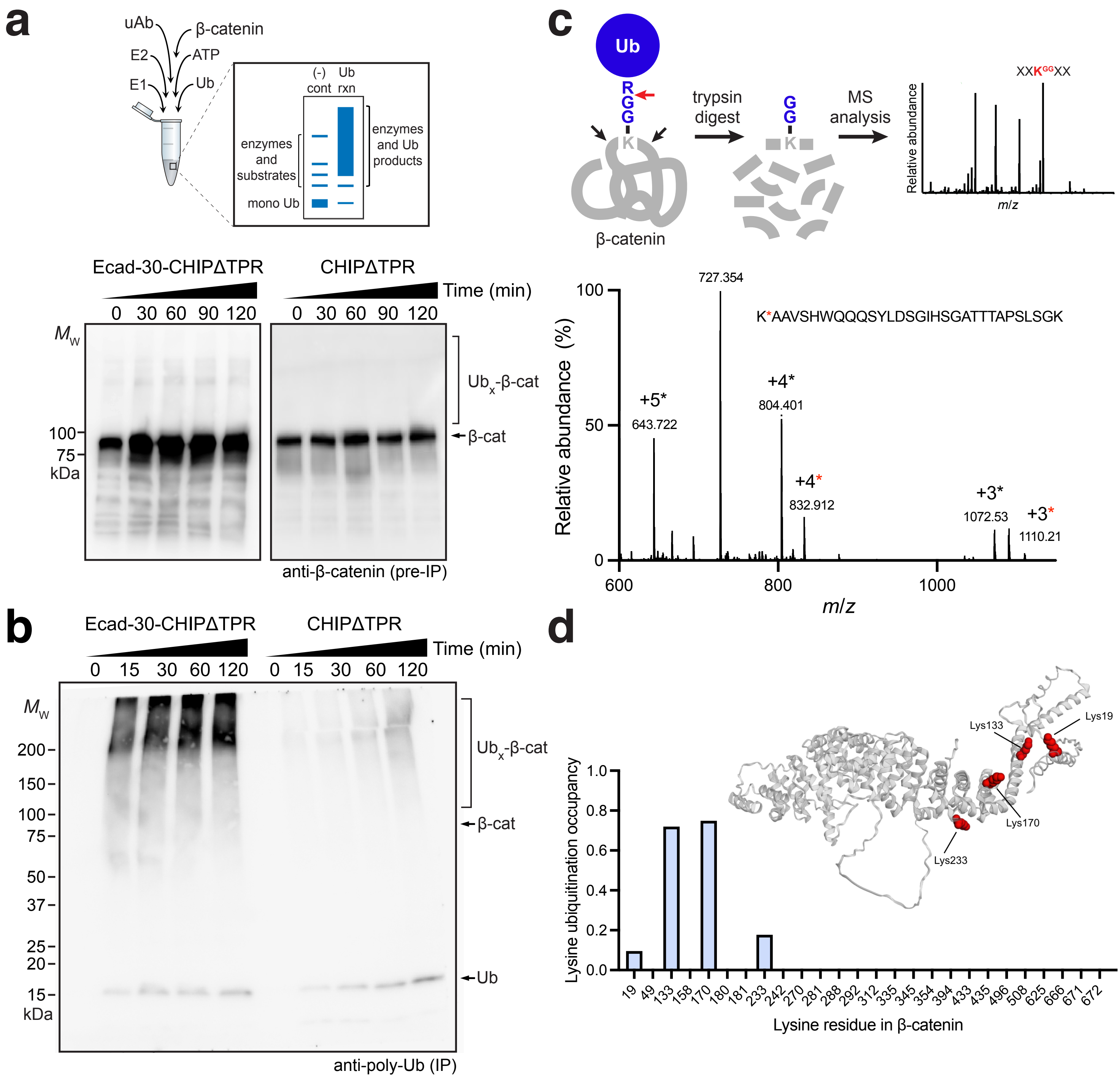


**Supplementary Figure 6. *In vitro* ubiquitination of β-catenin by peptide-guided uAb.** (a) *In vitro* ubiquitination of β-catenin in the presence of purified Ecad-30-CHIPΔTPR or CHIPΔTPR along with E1, E2, ubiquitin (Ub), and ATP. Samples were collected at indicated times and subjected to immunoblotting with anti-β-catenin antibody. The figure was made using BioRender.com. (b) Immunoblot analysis of samples in (a) following immunoprecipitation (IP) using beads coated with β-catenin-specific VHH, BC13. Lanes were normalized by total protein content and molecular weight ($M_W$) markers are indicated at left. All blots are representative of biological replicates ($n$ = 3). (c) Mass spectrometry (MS)-based identification of ubiquitin attachment sites based on characteristic mass shift caused by presence of diglycine (GG) that is retained on ubiquitinated lysine residues within peptides after trypsin digestion. Raw ESI-MS/MS data for K19 polyubiquitinated β-catenin peptide with each number marking a different charged state, signals corresponding to the unmodified peptide are marked with a black asterisk and those corresponding to the ubiquitinated peptide are marked with a red asterisk. (d) Lysine occupancy rate based on GG modification of β-catenin as determined by LC-MS/MS. Peptides corresponding to 80% of the β-catenin sequences were identified using Mascot software. Data were generated by normalizing ubiquitinated residue counts relative to total residue counts and by averaging across three independent experiments. Ubiquitination sites are shown in red in the structure of the AlphaFold2-predicted β-catenin structure.

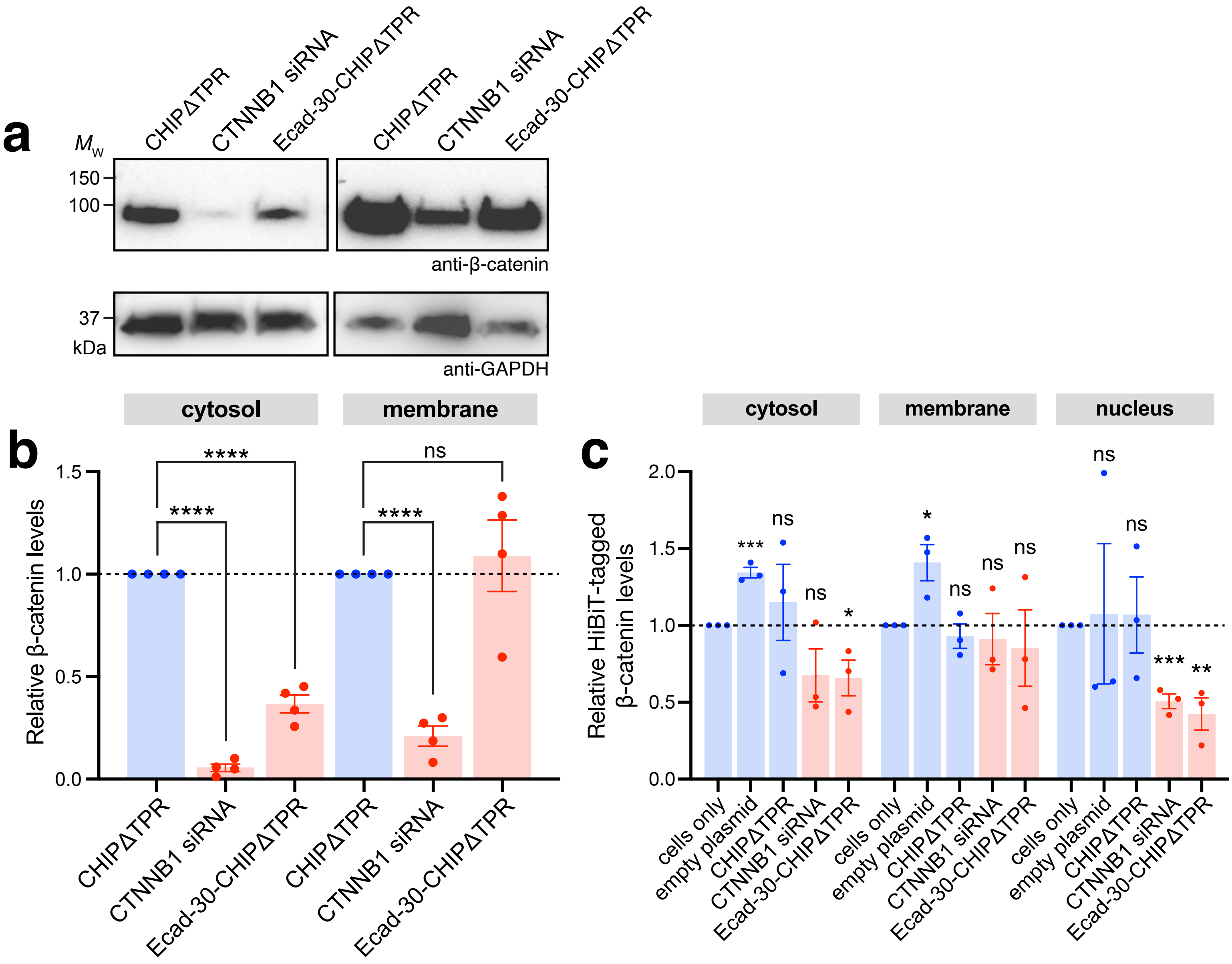


**Supplementary Figure 7. Comparison of β-catenin knockdown by peptide-guided uAb versus siRNA.** (a) Immunoblot analysis of cytosolic and membrane β-catenin levels in DLD1 cells transfected with either β-catenin-specific siRNA or plasmid pcDNA3 encoding CHIPΔTPR or Ecad-30-CHIPΔTPR. Cells were harvested 48 h post-transfection, after which cytoplasmic and membrane fractions were prepared from cell extracts and subjected to immunoblotting with anti-β-catenin antibody (top) and anti-GAPDH antibody (bottom), the latter serving as a loading control for both cytosolic and membrane fractions. Lanes were normalized by total protein content and molecular weight ($M_w$) markers are indicated at left. Blots are representative of four biological replicates. (b) Quantification of cytosolic and membrane β-catenin levels by densitometry analysis of immunoblots in panel (a). Band intensity was determined using ImageJ software with all β-catenin band intensities normalized to corresponding GAPDH band intensities. Relative β-catenin levels were then calculated by normalizing Ecad-30-CHIPΔTPR values to CHIPΔTPR values. Data are mean of biological replicates ($n$ = 4) ± SD. Statistical significance was determined by unpaired two-tailed Student's $t$-test. Calculated $p$ values are represented as follows: *, $p < 0.05$; **, $p < 0.01$; ***, $p < 0.001$; ****, $p < 0.0001$; ns, not significant.

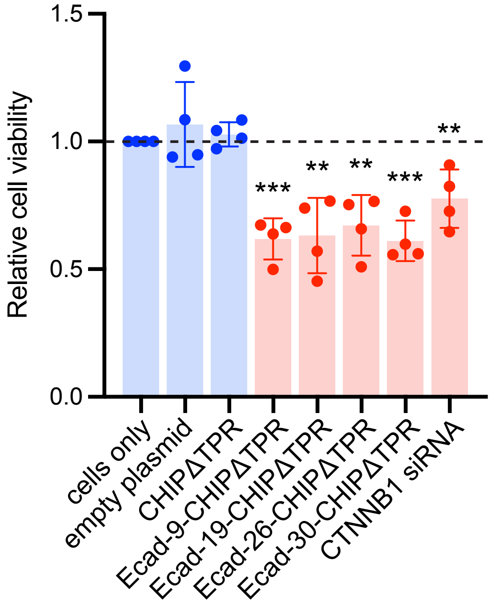


**Supplementary Figure 8. Effect of peptide-guided uAb degraders on cell viability.** Viability of DLD1 cells transfected with empty plasmid pcDNA3, pcDNA3 encoding CHIPΔTPR or one of the peptide-guided uAb degraders, or β-catenin-specific siRNA. Cells were harvested 24 h post-transfection, after which viability was quantified by MTS assay. Data are mean of biological replicates ($n$ = 4) ± SD. Statistical significance in all panels was determined by unpaired two-tailed Student's $t$-test. Calculated $p$ values are represented as follows: **, $p < 0.01$; ***, $p < 0.001$.

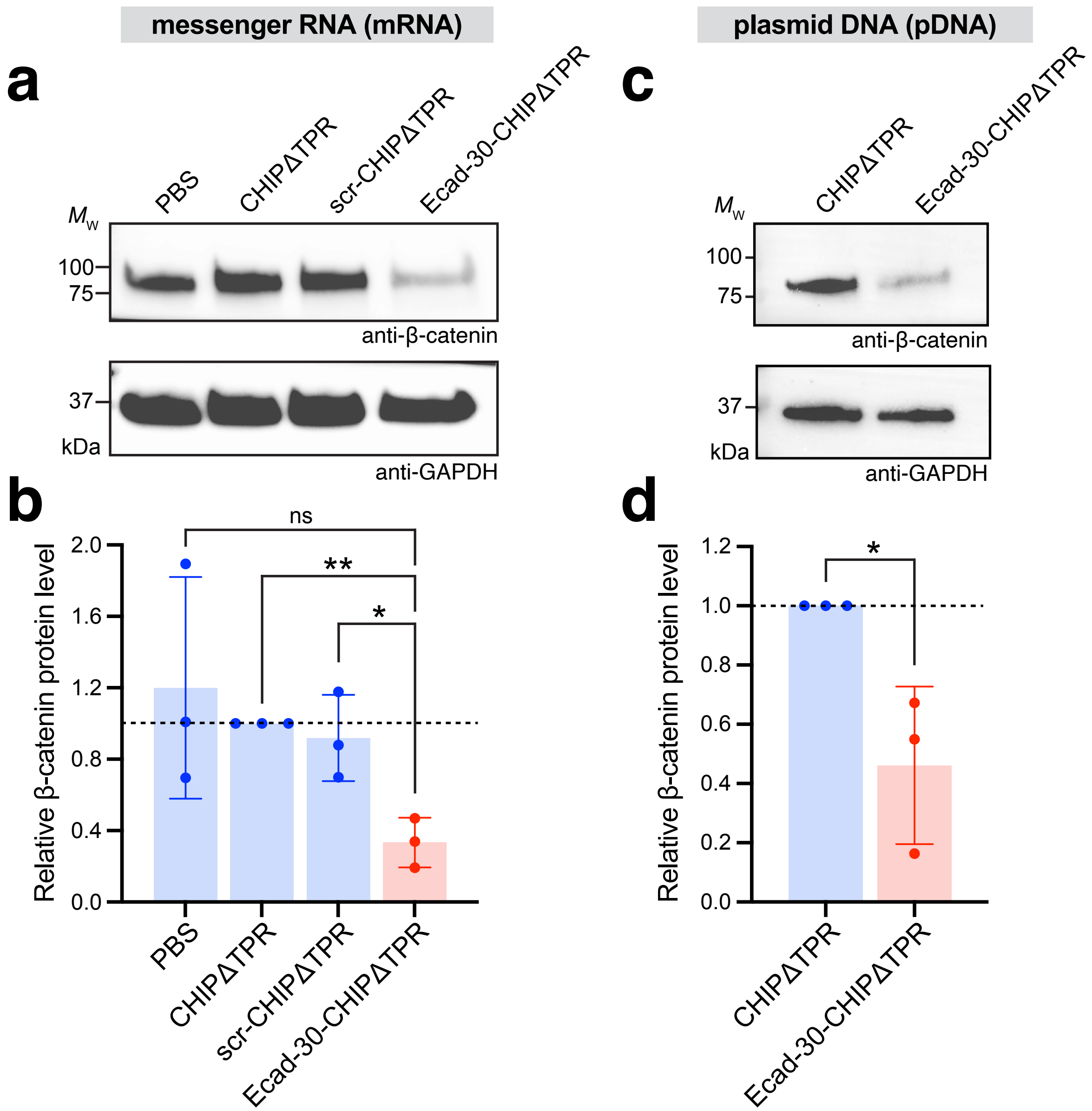


**Supplementary Figure 9. Cytosolic β-catenin knockdown by pDNA and mRNA in cultured Hep3B cells.** (a) Immunoblot analysis of cytosolic β-catenin levels in Hep3B cells transfected with LNP-encapsulated mRNA encoding CHIPΔTPR, scr-CHIPΔTPR, or Ecad-30-CHIPΔTPR as indicated. Treatment of cells with PBS served as a negative control. Cells were harvested 48 h post-transfection, after which cytoplasmic fractions were prepared from cell extracts and subjected to immunoblotting with anti-β-catenin antibody (top) and anti-GAPDH antibody (bottom), the latter serving as a loading control. Lanes were normalized by total protein content and molecular weight ($M_W$) markers are indicated at left. Blots are representative of at least three biological replicates. (b) Quantification of cytosolic β-catenin levels by densitometry analysis of immunoblots in panel (a). Band intensity was determined using ImageJ software with all β-catenin band intensities normalized to corresponding GAPDH band intensities. Relative β-catenin levels were then calculated by normalizing all values to CHIPΔTPR values. Data are mean of biological replicates ($n$ = 3) ± SD. (c, d) Same as (a,b) but Hep3B cells were transfected with plasmid pcDNA3 encoding CHIP ΔTPR or Ecad-30-CHIPΔTPR as indicated. Statistical significance was determined by unpaired two-tailed Student's $t$-test. Calculated $p$ values are represented as follows: *, $p < 0.05$; **, $p < 0.01$; ns, not significant.

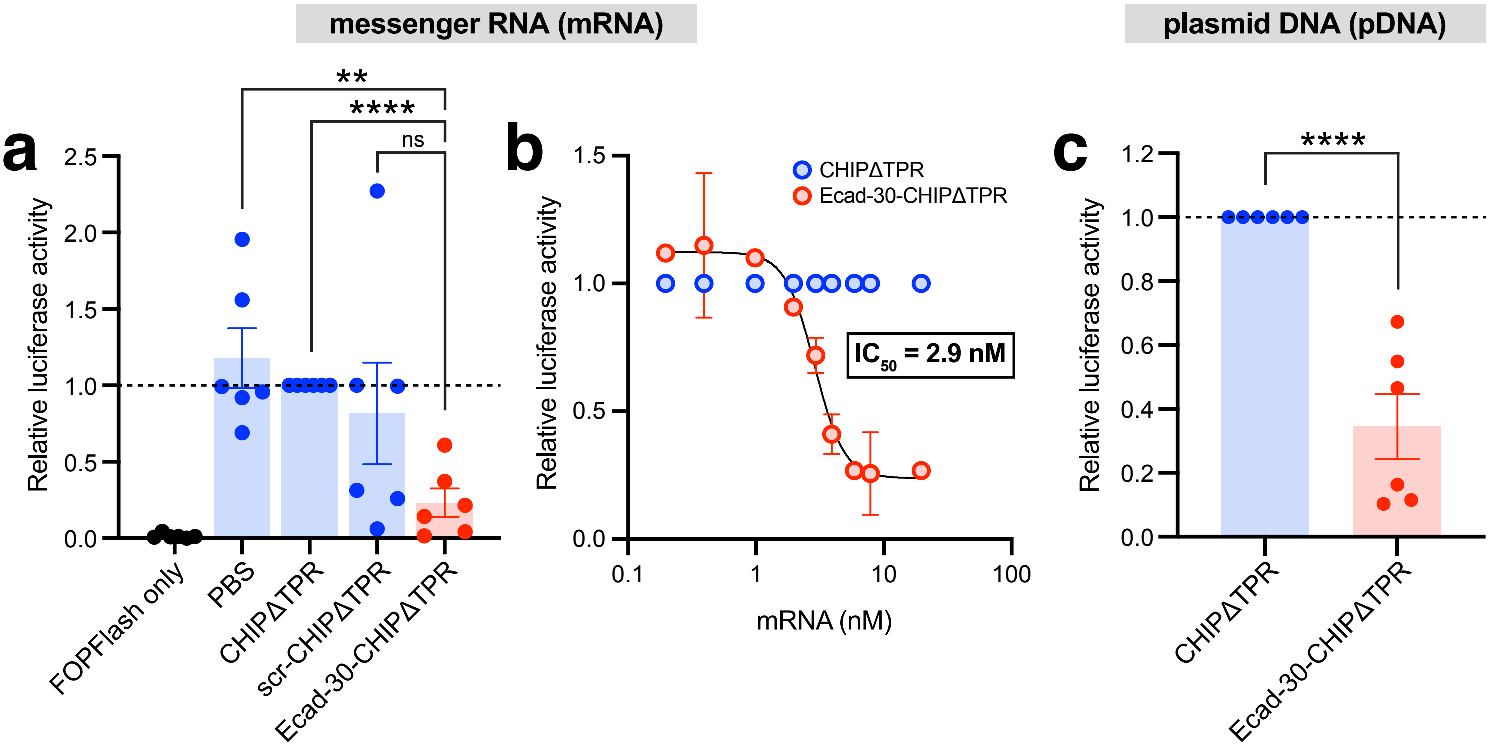


**Supplementary Figure 10. Inhibition of β-catenin signaling by pDNA and mRNA in cultured Hep3B cells.** (a) β-catenin signaling activity in Hep3B cells co-transfected with TOPFlash reporter plasmid along with LNP-encapsulated mRNA (7.86 nM) encoding CHIPΔTPR, scr-CHIPΔTPR, or Ecad-30-CHIPΔTPR as indicated. Treatment of cells with PBS served as a negative control. Cells receiving only the FOPFlash plasmid served as additional negative control. Cells were harvested 48 h post-transfection. Luciferase signals in each sample were normalized to those measured in cells receiving LNP-encapsulated mRNA encoding CHIPΔTPR. Data are mean of biological replicates ($n$ = 6) ± SD. (b) β-catenin signaling activity in Hep3B cells receiving varying concentations (0.2–20 nM) of LNP-encapsulated mRNA encoding CHIPΔTPR or Ecad-30-CHIPΔTPR. Data are the mean of biological replicates ($n$ = 3) ± SD. (c) Same as in (a) but Hep3B cells were transfected with plasmid pcDNA3 encoding CHIPΔTPR or Ecad-30-CHIPΔTPR as indicated. Data are mean of biological replicates ($n$ = 6) ± SD. Statistical significance was determined by unpaired two-tailed Student's $t$-test. Calculated $p$ values are represented as follows: **, $p < 0.01$; ****, $p < 0.0001$; ns, not significant.

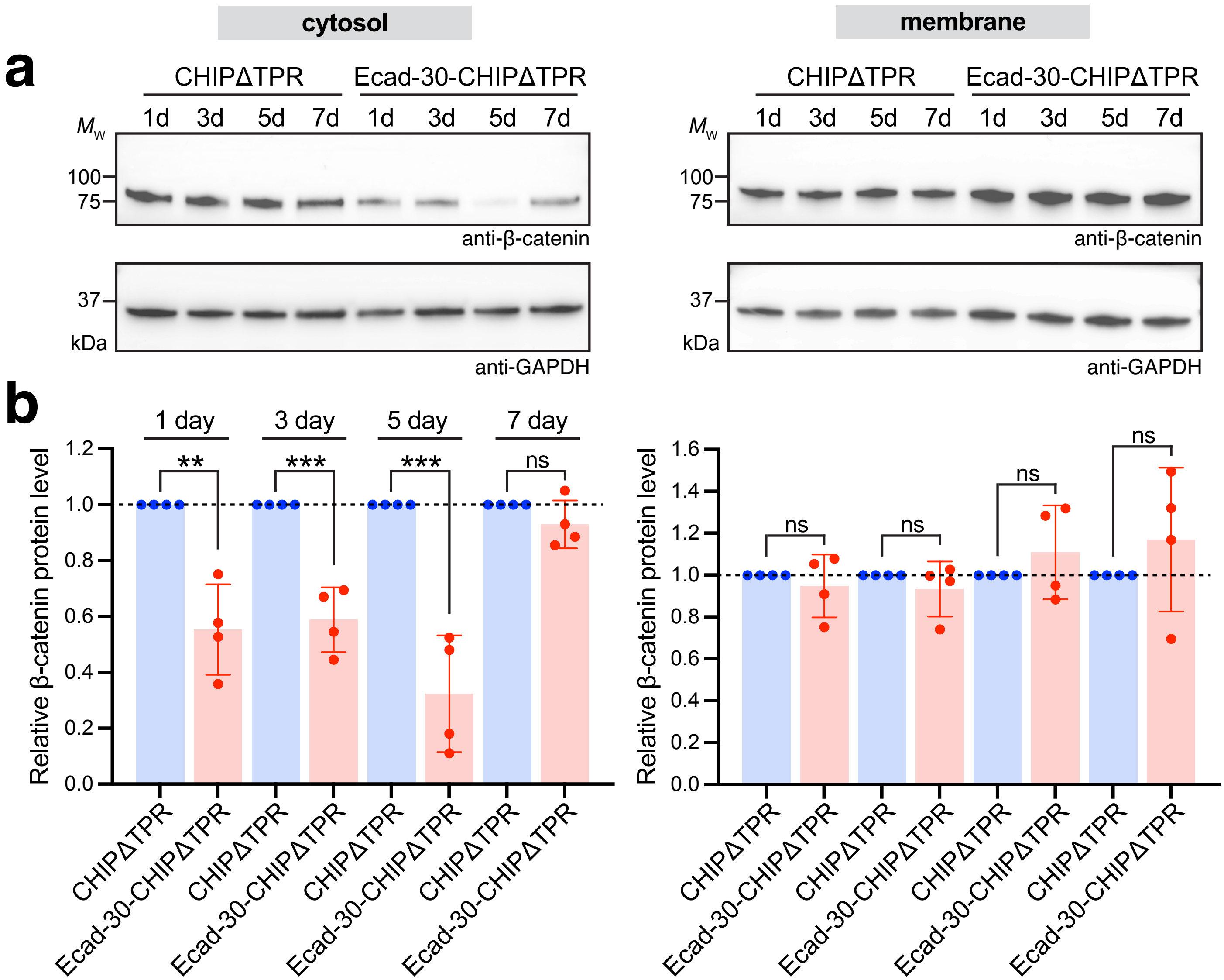


**Supplementary Figure 11. Silencing of cytosolic β-catenin following LNP-mediated delivery of uAb mRNA in mice.** (a) Representative immunoblot analysis of β-catenin in cytosolic (left) and membrane (right) fractions derived from homogenized livers of wild-type BALB/c mice ($n$ = 4 mice per group). Mice were i.v. injected with a single 1.0 mg/kg dose of the following: PBS, CHIPΔTPR-mRNA-LNP, scr-CHIPΔTPR-mRNA-LNP, or Ecad-30-CHIPΔTPR-mRNA_LNP and livers were collected at 1, 3, 5 and 7 days post injection. Blots were probed with anti-β-catenin antibody (top) and anti-GAPDH antibody (bottom), the latter serving as a loading control for both cytosolic and membrane fractions. Lanes were normalized to the tissue weight and total protein for each liver. Molecular weight ($M_w$) markers are indicated at left. Blots are representative of two technical replicates per mouse. (b) Quantification of cytosolic and membrane β-catenin levels by densitometry analysis of immunoblots. Band intensity was determined using ImageJ software with all β-catenin band intensities normalized to corresponding GAPDH band intensities. Relative β-catenin levels were then calculated by normalizing values to CHIPΔTPR control. Data are mean of biological replicates ± SD, where each data point is the average of two technical replicates. Statistical significance was determined by unpaired two-tailed Student's $t$-test. Calculated $p$ values are represented as follows: **, $p < 0.01$; ***, $p < 0.001$; ns, not significant.
